# Loss of Hem1 disrupts macrophage function and impacts on migration, phagocytosis and integrin-mediated adhesion

**DOI:** 10.1101/2020.03.24.005835

**Authors:** Stephanie Stahnke, Hermann Döring, Charly Kusch, David J.J. de Gorter, Sebastian Dütting, Aleks Guledani, Irina Pleines, Michael Schnoor, Michael Sixt, Robert Geffers, Manfred Rohde, Mathias Müsken, Frieda Kage, Anika Steffen, Jan Faix, Bernhard Nieswandt, Klemens Rottner, T.E.B. Stradal

## Abstract

The hematopoietic-specific protein 1 (Hem1) comprises an essential subunit of the WAVE Regulatory Complex (WRC) in immune cells. WRC has a fundamental role in Arp2/3 complex activation and the protrusion of branched actin networks in motile cells.

Hem1 deficiency leads to suppression of the entire WRC in immune cells. Defective WRC function in macrophages results in loss of lamellipodia and migration defects. Moreover, phagocytosis, commonly accompanied by lamellipodium protrusion during cup formation, is altered in Hem1 null cells concerning frequency and efficacy. When analyzing cell spreading, adhesion and podosome formation, we found that Hem1 null cells are capable, in principle, of podosome formation and consequently, do not show any quantitative differences in extracellular matrix degradation. Their adhesive behavior, however, was significantly altered. Specifically, adhesion as well as de-adhesion of Hem1 null cells was strongly compromised, likely contributing to the observed reduced efficiency of phagocytosis. In line with this, phosphorylation of the prominent adhesion component paxillin was diminished. Non-hematopoietic somatic cells disrupted in expression for both Hem1 and its ubiquitous orthologue Nck-associated protein 1 (Nap1) or the essential WRC components Sra-1/PIR121 did not only confirm defective paxillin phosphorylation, but also revealed that paxillin turnover in focal adhesions is accelerated in the absence of WRC. Finally, adhesion assays using platelets lacking functional WRC as model system unmasked radically decreased αIIbβ3 integrin activation.

Our results thus demonstrate that WRC-driven actin networks impact on integrin-dependent processes controlling formation and dismantling of different types of cell-substratum adhesion.

**One sentence summary:** Interference of Hem1 function in mice and cells uncovers a hitherto unrecognized role in integrin-mediated cell adhesion that is crucial for macrophage function and connects to recently discovered immunodeficiencies in patients carrying Hem1 mutations.

## INTRODUCTION

Cell migration follows an orchestrated sequence of events, a cycle of cell protrusion, adhesion to newly occupied space and contraction to drag the cell body forward. Polymerization of actin filaments (F-actin) at the cell front pushes the plasma membrane into the direction of migration, which then attaches to the extracellular substratum via members of the integrin transmembrane receptors family. Cells in metazoan organisms are capable of migrating through complex, 3-dimensional environments and their license keys to different organismal compartments constitute combinations of integrins they carry on their surface, allowing them to specifically adhere to given substrata (*1, 2*). Once engaged, the integrins dynamically couple to the actin cytoskeleton, which in turn can exert pulling forces through myosin II activation and contraction, culminating in locomotion of the cell body (*3-5*)

Integrins are heterodimeric cell surface receptors that are responsible for cell adhesion during several biological processes. Lammermann and colleagues (*6*) have shown that dendritic cells (DCs) are capable of migrating by solely employing the forces derived from actin-network expansion and protrusive flowing of the leading edge. Myosin II-dependent contraction is only required for squeezing through narrow pores (*6, 7*). This mode of movement, however, only works for 2D migration or in 3D through loose mazes, and fails when cells are transmigrating through tight tissue barriers, as this requires the ‘license key’ of integrin-mediated adhesion (*6, 8*). More recently, we analyzed DC migration in the absence of WRC-dependent lamellipodium formation (*9*). WRC is a hetero-pentameric complex, mediating lamellipodium protrusion through Arp2/3 complex activation and formation of the branched lamellipodial actin network that pushes the membrane forward (*10, 11*). WRC was inactivated by interfering with expression of its hematopoietic subunit Hem1, which causes loss of function and suppression of expression of essential other WRC subunits, such as WAVE (*9, 12-14*). Remarkably, WRC-deficient cells were observed to migrate in 2D with increased speed, but were ineffective at turning to follow chemotactic gradients. Moreover, their migration was progressively impaired with increasing geometrical complexity of the extracellular environment (*9*). Thus, diversified leading edge protrusions enable leukocytes to sense and explore their environment, whereas integrin-mediated adhesion is required for switchable immobilization to specifically assigned surfaces leading to retention, invasion, or cell-cell communication (*7, 9, 15*).

Studying DC migration has allowed to dissect these two processes, WRC-mediated cell edge protrusion *versus* integrin-mediated adhesion. However, studies in lymphocytes are revealing that these events may also depend on each other: In case of T-cells, conjugation to an antigen presenting cell (APC) or target cell leads to cell-cell-adhesion and formation of the immunological synapse (IS). T-cells express different integrins, of which leukocyte-function-associated-(LFA)-1 (α_L_β_2_) is known to be most critical for conjugate formation as it binds to its ligand ICAM-1 on APCs (*16*). Actin rearrangements that occur concomitantly with IS formation and T-cell activation are well established to involve WRC (*17-19*). Moreover, WRC mediates integrin affinity maturation within the IS by recruiting vinculin and talin in an Arp2/3 complex-dependent manner. Finally, Noltz and colleagues (*20*) showed that the Abl tyrosine kinase associates with WRC upon T-cell-receptor (TCR)-ligation at the nascent IS, leading to Abl-mediated Rap1-activation in a WRC-Abl-CrkL–C3G dependent fashion (*20, 21*). Hence, during IS formation, WRC is required for TCR-mediated activation of the integrin regulatory small GTPase Rap1 and thus integrin activation.

To shed more light on the role of WRC-driven Arp2/3 complex-mediated protrusion and the intricate interplay with contraction and adhesion in motile cells, we chose to analyze the motility of macrophages derived from mice genetically deficient for Hem1. We then compared our results with those from other cell types, namely platelets from the same mouse model and from CRISPR/Cas9 genome-edited cancer cell lines. WAS protein (WASP) embodies the second exclusively hematopoietic activator of Arp2/3-complex and mutations are causative of the well characterized X-linked immune deficiency Wiskott Aldrich Syndrome (*22*) In order to demarcate WASP-phenotypes from WRC loss of function, we included cells from mice lacking WASP (*23*) in our analyses.

Our results demonstrate for the first time that WRC has a specific and conserved function in the regulation of cell adhesion during motile processes in addition to its well-accepted role in the formation of lamellipodial protrusions. This highlights WRC as a master conductor in orchestrating the halting and going of cells, independent of the specific integrin engaged.

## RESULTS

### 1. Cell morphology, migration and polarization of Hem1-null macrophages

Macrophages when allowed to spread on fibronectin assume a flat shape with smooth edges. During this process they form focal adhesions and, depending on their state of differentiation and activation, potentially also podosomes. Spreading of macrophages on fibronectin is accompanied by both WRC-mediated lamellipodium protrusion and engagement of the integrin LFA-1 (αLβ2). *In vitro*-differentiated macrophages were derived from the bone marrows of WT and Hem1 knockout C57BL/6 mice, being WRC defective in the latter case (*9*), or from mice lacking WASP (*23*). Whereas macrophages of all three genotypes readily adhered to fibronectin, WRC-deficient *hem1* null macrophages clearly showed an aberrant, spiky cell shape with multiple filopodia (Fig. 1A, Movie S1A). WT and WASP-null cells did not display obvious differences in these conditions (Fig. 1A). To probe whether the loss of WASP or WRC expression affects expression levels of the respective other Arp2/3 complex activator, we performed immunoblotting. No compensatory expression of either WRC or WASP was observed in the respective other knockout (Figs. 1B and S1A). Moreover, Hem1 deletion leads to significant loss of WAVE and the other WRC components – as expected from previous work (compare (*9, 13, 14*)). The Abl interacting protein Abi-1 was also significantly reduced (Fig. 1B, Fig. S1B), but as it is believed to also participate in other complexes (*24-26*), we probed localization of the remaining Abi-1 protein. Whereas Abi-1, as expected for a WRC component, readily localizes to the plasma membrane and accumulates at lamellipodia tips and ruffles of WT macrophages, these staining characteristics are lacking in Hem1 KO cells, with the signal being diffusely distributed throughout the cytoplasm (Fig. 1C).

**Figure 1.**
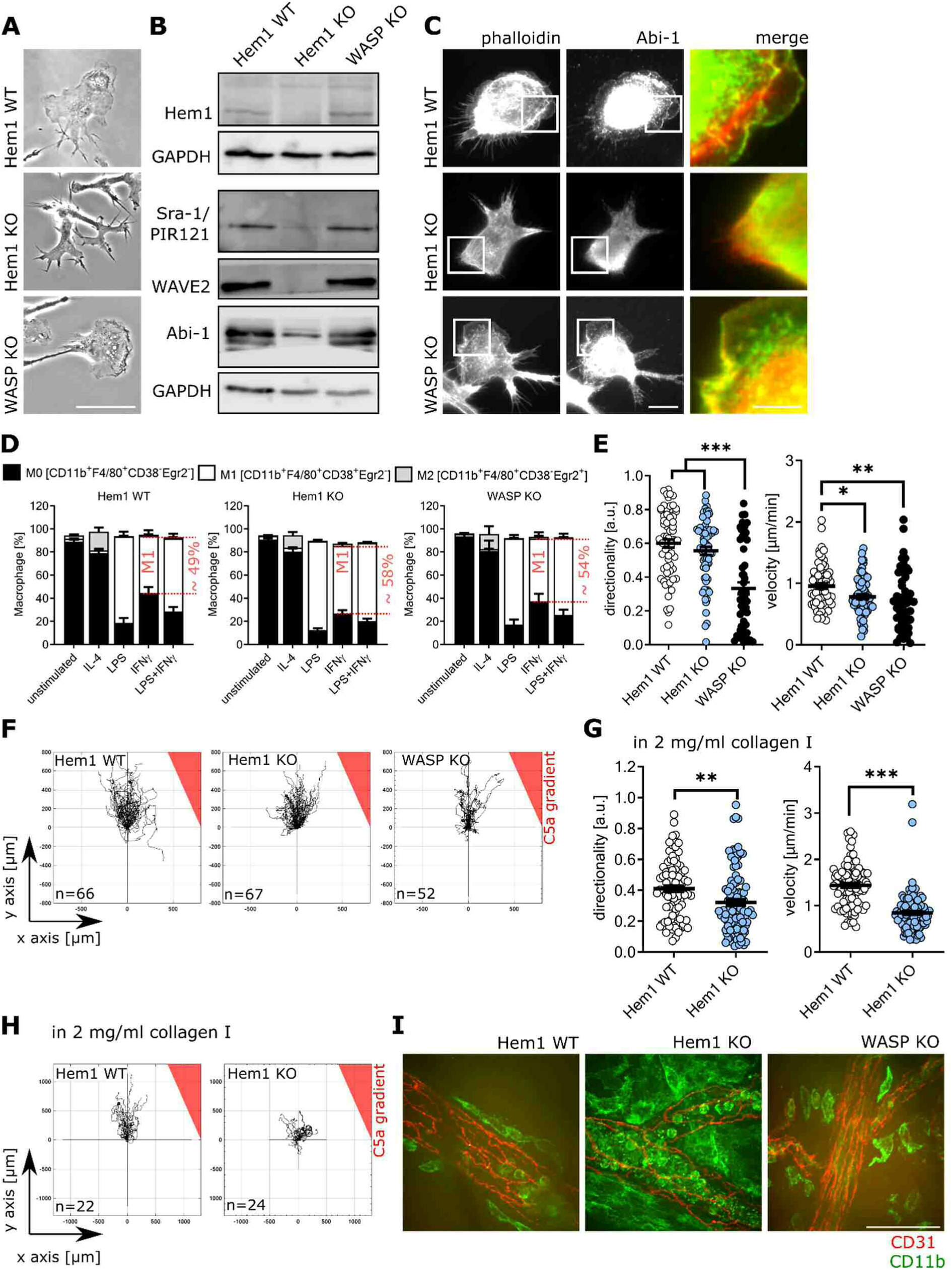
Differentiation and migratory performance of Hem1 null macrophages. **(A)** Still images from time-lapse videos illustrating the lack of ruffles and lamellipodia in Hem1 KO macrophages. Bar=50µm, in merge bar=5µm (n=3). **(B)** Representative Western blots demonstrating virtual loss of WRC components Sra-1 and WAV2, and at least partial reduction of Abi-1 protein levels in Hem1 KO macrophages. Loading control is GAPDH (n=5). **(C)** Peritoneal macrophages stained with phalloidin and α-Abi-1, revealing the absence of leading edge localization of Abi-1 upon Hem1 but not WASP knockout. Bar=10µm. **(D)** Macrophage polarization status in WT, Hem1 KO and WASP KO cells as indicated, identified via assessment of expression of Egr2 (for M2) or CD38 (for M1) after 24h stimulation with indicated stimulants. Absence of either marker was defined as M0 state. Respective percentage of M1 for IFNγ is indicated in red (n=3; 3 replicates). **(E)** Quantification of directionality (left side) and velocity (right side) from a 2D migration assay of macrophages crawling towards a C5α gradient (n=3). In total 66 WT, 67 Hem1 KO and 52 WASP KO macrophages were analyzed. **(F)** Trackplots for the 2D migration data quantified in (E). **(G)** 3D migration assay of Hem1 KO and WT macrophages in a collagen I matrix (2 mg/ml) towards a C5α gradient. Quantification of directionality (left side) and velocity (right side). In total, 89 WT and 104 Hem1 KO macrophages were analyzed. (H) Trackplots for the 3D migration assay quantified in (G). **(I)** Immunofluorescence staining of cremaster muscles of male mice from the three genotypes as indicated. Vessel borders are stained for endothelial cell contact protein CD31 (red) and the myeloid cell integrin component CD11b (green). Bar=50µm.

We next asked whether the deviating cell morphology of *in vitro*-differentiated macrophages might be due to a differentiation defect. Therefore, we probed expression of the markers CD11b and ADGRE (also known as F4/80), the combination of which identifies murine macrophages, and found no differences (Fig. S1C). Next, we analyzed the polarization towards M1/M2 of *in vitro*-differentiated macrophages of all three genotypes by flow cytometry upon different stimulations in more detail. IL-4 is known to induce alternative activation of macrophages leading to so called wound healing or M2 macrophages (*27*), characterized by Erg2 expression, while danger signals such as interferon gamma (IFNγ) and bacterial LPS induce the classical activation of macrophages towards the inflammatory M1 phenotype (*28*), characterized by markers such as CD38 (*29*). No dramatic changes in the polarization of bone marrow-derived macrophages could be detected, although there was a trend towards a higher percentage of M1 macrophages in Hem1 null cells upon IFNγ stimulation (Fig. 1D, S1D).

To understand if or to which extend the loss of individual Arp2/3 complex activators affects the migratory performance of macrophages, we performed chemotaxis assays towards C5a on 2D surfaces (Fig. 1E-F,), and in 3D collagen matrices (Fig. 1 G-H). In earlier studies, we had analyzed the migration of dendritic cells (DCs) from Hem1 KO mice (*7, 9*), which showed that Hem1 null DCs formed unipolar cells migrating with increased speed and enormous directional persistence on 2D surfaces, whereas they were unable to turn towards chemotactic gradients. In 3D environments, Hem1 null DCs failed to make directional decisions due to compromised substrate exploration (*9*). However, as opposed to the integrin-independent migration of DCs (*6*), macrophages assume a migration mechanism employing excessive spreading and different types of integrin-mediated adhesions. Hem1 null macrophages were capable to chemotax under all conditions, but were slightly reduced in their velocity (Fig. 1E-F). In line with the literature, WASP KO cells displayed stronger chemotactic defects (*30-36*). However, when migrating in 3D collagen matrices, Hem1 null macrophages displayed significant defects concerning both their velocity and directionality (Fig. 1 G, H), whereas in 2D chemotaxis, these parameters only displayed a slight trend towards reduction (Fig. 1E, F). Together, the migratory performance of macrophages lacking Hem1 and thus being incapable of canonical lamellipodia formation displayed a specific phenotype, different from that seen in either DCs lacking Hem1 (*9*) or WASP KO macrophages (*32*). In order to learn whether this phenotype is also reflected in changes of myeloid migration *in vivo*, we turned to the analysis of leukocyte extravasation in cremaster muscle. Strikingly, blood vessels from mice lacking Hem1, but not WT or WASP KOs, were prominently filled with myeloid cells. Moreover, and despite a potentially reduced capacity to migrate in 3D or to extravasate, the interstitial space around the vessels was overly populated with cells that have likely accumulated over time (Fig. 1I, Movie S1B). This extreme phenotype is reminiscent of an acute, local inflammation (*37*) and indicative of deviant myeloid extravasation cascade and/or motility following extravasation.

### 2. Podosome formation, extracellular matrix degradation and adhesion of WRC-deficient macrophages and somatic cells

A very important feature of motile behavior in general and macrophage function in particular is integrin mediated adhesion, which is involved in cell-substrate and cell-cell adhesion as well as in phagocytosis (*38-40*). Macrophages also form specialized types of cell-substratum adhesion, termed podosomes. These structures constitute dot-like matrix contacts that structurally and functionally differ from cell-substrate adhesions in non-hematopoietic cells, but are related to invadopodia in cancerous cells (*41, 42*). Structurally, they show a bipartite architecture with a core of Arp2/3 complex-nucleated F-actin and actin-associated proteins, surrounded by a ring consisting of adhesion proteins like vinculin (*43*). Moreover, the ability of podosomes and invadopodia to engage in matrix degradation sets them functionally apart from other cell-matrix contacts. It is commonly agreed that WASP constitutes the major activator of Arp2/3 complex in the podosome actin core. However, it has remained unclear whether the ability to form lamellipodial protrusions is involved in this process at any stage. Therefore, we analyzed podosome formation in macrophages of all three cell types, Hem1 KO, WT and WASP KO. Peritoneal macrophages isolated from WT mice form podosomes on gelatin, characterized by the typical actin rich core surrounded by an adhesive, vinculin-containing ring. The same structures were readily observed in Hem1 KO macrophages. In contrast, WASP KO macrophages only very rarely displayed podosomes (Fig. 2A). This is in line with work identifying WASP as essential component of podosome formation (*32*), which only occasionally can be substituted by penetrating expression of the non-hematopoietic isogene N-WASP (*36*). Next, we asked whether the podosomal activity of extracellular matrix degradation might be altered in the absence of Hem1. To do so, we seeded macrophages of all three genotypes onto fluorescent gelatin and assessed the degraded area, identified by loss of the fluorescent signal after 20 hours. Interestingly, both WT and Hem1 KO macrophages readily degraded the extracellular matrix beneath them, coinciding with podosome formation. These typical areas of gelatin degradation as well as podosomes were absent from macrophages lacking WASP (Fig. 2B). Occasionally gelatin was also degraded beneath WASP KO macrophages, which then however coincided with ventral ruffling of respective cells (not shown), indicating that podosomes may not be absolutely required for gelatin degradation. Quantification of the area of gelatin removal (Fig. 2C) confirmed that there is no detectable difference between the degradation efficacies of WT- and Hem1-KO-macrophages, and underscored the significant defect in WASP KO cells.

**Figure 2.**
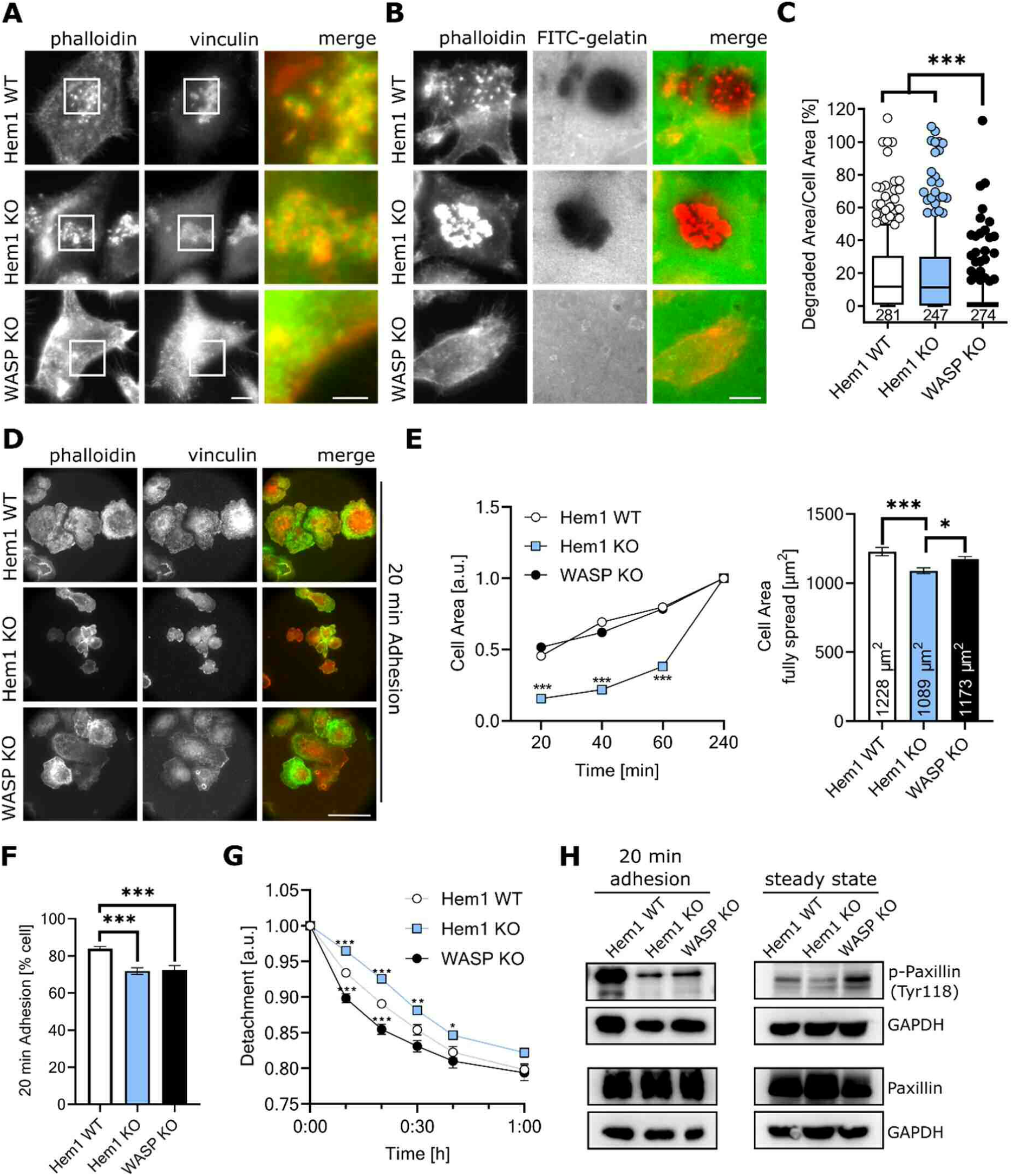
Adhesion characteristics and podosome functionality in Hem1 null macrophages. **(A)** Vinculin and phalloidin staining of peritoneal macrophages to detect podosomes in the three genotypes. Note that WASP KO cells are defective for these structures. Bar=10µm, in merge bar=5µm. **(B)** FITC-gelatin degradation assay. Note that degradation appears as dark area beneath WT- and Hem1 KO, but not WASP KO macrophages. Bar=10µm. **(C)** Quantification of degraded area per cell area as determined from images in (B). Data are displayed as median boxplots with 10-90 percentile whiskers from two independent experiments (n=2). Values below bars represent the total amount of cells measured. **(D)** Immunofluorescence of macrophages allowed to spread for 20min, and fixed and stained for vinculin (red) and F actin (green). Scale bar is 50µm. **(E)** Left: Quantification of the cell area of macrophages during spreading on fibronectin. Values are normalized to respective endpoint value for each cell type, to illustrate the delayed course of spreading in Hem1 null cells. Right: Mean area of macrophage genotypes after 4h spreading on fibronectin. The average value for each genotype is displayed within the respective bar. **(F)** Adhesion assay. Percentage of attached macrophages after 20min adhesion to fibronectin and washing off of non-adherent cells (n=3; 12 replicates). Note the reduction of app. 15% cells capable of adhesion in these conditions in the absence of both Hem1 and WASP. **(G)** Kinetics of detachment of macrophages as measured by decreasing eccentricity (n=3; 12 replicates), which is delayed in Hem1 null but not WASP KO cells. Values are normalized to respective starting point for each cell type. **(H)** Representative Western blots of paxillin and phospho-paxillin (p-Paxillin [Tyr118]) expression during steady state culture (16h after seeding) (n=3) and after 20min of cell adhesion (n=4) as depicted in (D).

During routine tissue culture, we noted that bone marrow-derived Hem1 KO macrophages spread significantly slower than WT cells 20 min after seeding (Fig. 2D). While this was evident for Hem1 KO cells, cell size of WASP KO cells appeared to be virtually identical to WT cells (Fig. 2D, E). Upon prolonged spreading times and in flow cytometry assays, no major difference in cell size was apparent for all genotypes. To further explore the efficiency of adhesion of our cell types, we next explored how many cells stayed attached to fibronectin following 20 minutes of spreading and washing. In fact, a lower number of Hem1 KO and WASP KO cells had effectively adhered as compared to WT macrophages (Fig. 2F). However, 2-3 hours later, all cells were fully spread, showing that cell adhesion is principally not defective, but delayed. We then asked if decelerated adhesion in the absence of WRC also results in less firm adhesion. To test this hypothesis, we analyzed the speed of cell rounding upon EDTA-mediated detachment. Despite delayed cell adhesion, Hem1 KO cells displayed significantly sustained adhesion once attached to the substratum, compared to WT. Interestingly, WASP KO macrophages showed the opposite behavior and adhered even less firmly than WT cells, displaying very fast rounding upon addition of EDTA (Fig. 2G, Movie S2). This behavior of Hem1 KO macrophages is in agreement with our observation that they detach less easily in regular tissue culture.

To uncover the molecular basis of this phenomenon, we first assessed the RNA expression profile of WT *versus* Hem1 KO macrophages (Figs. S2A, S2B, Table S1), followed by analysis of the expression of adhesion complex proteins (Fig. 2H, S2C-S2D) and finally their phosphorylation status (Fig. 2H, S2E-S2F). Pathway analyses of the expression profiles derived from Hem1 WT *versus* KO macrophages revealed that the RNA of the integrin alpha-subunits, of talin and vinculin, but not of paxillin or VASP are upregulated in Hem1 KO cells (Fig. S2B). We first probed if the adhesion components alpha-integrin and vinculin are also upregulated in expression at the protein level. Whereas surface expression levels of alpha-integrin, as determined using antibodies directed against CD11b (Integrin αM), were unaltered or slightly increased by trend, vinculin protein levels were reduced (Fig. S2C-D), indicating complex, additional layers of regulation in response to loss of WRC. Owing to the principal role of paxillin in signaling to adhesion formation and turnover (*44*), we further analyzed paxillin phosphorylation. Paxillin phosphorylation on tyrosine Y118 is induced upon integrin activation (*45*) and prominently involved in focal complex formation beneath lamellipodia as well as focal adhesion formation and turnover (*46, 47*). Western Blotting of paxillin confirms no significant regulation of its protein levels in Hem1 KO cells (Figs. 2H, S2E-F). Next, we monitored the tyrosine-phosphorylation status of paxillin in all three genotypes, WT, Hem1 KO and WASP KO. Strikingly, paxillin was found to be hypo-phosphorylated on residue Y118 both in spreading Hem1 KO cells (upon 20 minutes after seeding) and in steady-state adherent cells (16h after seeding) (Fig. 2H). Interestingly, WASP KO cells also displayed strongly reduced phosphorylation state during spreading (Fig. 2H, left panel), although this was even slightly increased as compared to WT at steady-state (Fig. 2H, right panel). Thus, phosphorylation of paxillin on Y118 negatively correlates with the detachment behavior/adhesion strength of all three genotypes tested, with low Y118 phosphorylation being associated with sustained adhesion and high levels of phospho-Y118 being linked to less firm adhesion (Fig. 2G-H). To better understand if the turnover of adhesion components is altered in the absence of functional WRC, we turned to cells that readily display canonical focal adhesions and can be easily transfected with tagged constructs.

We recently described B16-F1 melanoma cell lines consecutively depleted by CRISPR/Cas9-mediated genome editing for the ubiquitous Hem1 paralog and WRC-subunit Nap1 (*48*), followed by deletion of Hem1, giving rise to somatic, non-hematopoietic cell clones lacking functional WRC entirely. Moreover, we had previously characterized B16-F1 cells genetically inactivated for both *cyfip1* and *-2* genes, encoding the Rac GTPase-binding Sra-1 and its orthologue Pir121 (*49*). B16-F1 cells are a well-established model for lamellipodium formation, cell migration and adhesion dynamics (*48-52*). We here utilized these cells to address whether the impact of WRC loss of function on adhesion is restricted to macrophages or a more general phenomenon. Indeed, delayed initial adhesion (Fig. 3A-C) and sustained adhesion (Fig. 3D, Movie S3A) as well as reduced paxillin phosphorylation (Fig. 3E) were also detectable in different kinds of WRC-deleted B16-F1 cells. Moreover, reduction of vinculin expression was also mirrored in B16-F1 cells lacking WRC, although less pronounced than in macrophages (Fig. S3A). Thus, we conclude that B16-F1 cells are feasible for analyzing adhesion dynamics in more detail, specifically the turnover of individual focal adhesion molecules within these structures. Sra-1/PIR121 KO cells were transfected with either EGFP-tagged paxillin or – VASP, and turnover of these adhesion components assessed in fluorescence-recovery-after photobleaching (FRAP) experiments (Movies S3B and S3C). For both proteins VASP and paxillin, the mobile fraction (MF), reflecting the amount of protein that can be exchanged within given observation time, was slightly increased (Fig. 3F and S3B, raw data left graph), indicating that more of the respective protein is exchanging on average in focal adhesions of WRC-KO cells as compared to WT. As commonly observed for focal adhesion components, recovery curves do not display simple exponential fits to maximum, but follow bi-exponential functions, revealing at least two different sub-populations of molecules that recover with different velocities and individual half times of recovery, mirroring differentially phosphorylated subsets perhaps in case of paxillin. Indeed, phosphorylation of paxillin is known to affect adhesion formation and turnover (*53, 54*). However, the turnover rates of paxillin in the absence of WRC was increased for both subpopulations of molecules (slowly and rapidly cycling). In contrast, these values were unchanged for VASP (Fig. 3F, and S3B, fitted data right graph and Movie S3C). Together, formation and turnover of focal adhesions is significantly delayed in the absence of WRC, which is accompanied by reduced paxillin phosphorylation as well as accelerated paxillin turnover displaying a larger mobile fraction within individual adhesions.

**Figure 3.**
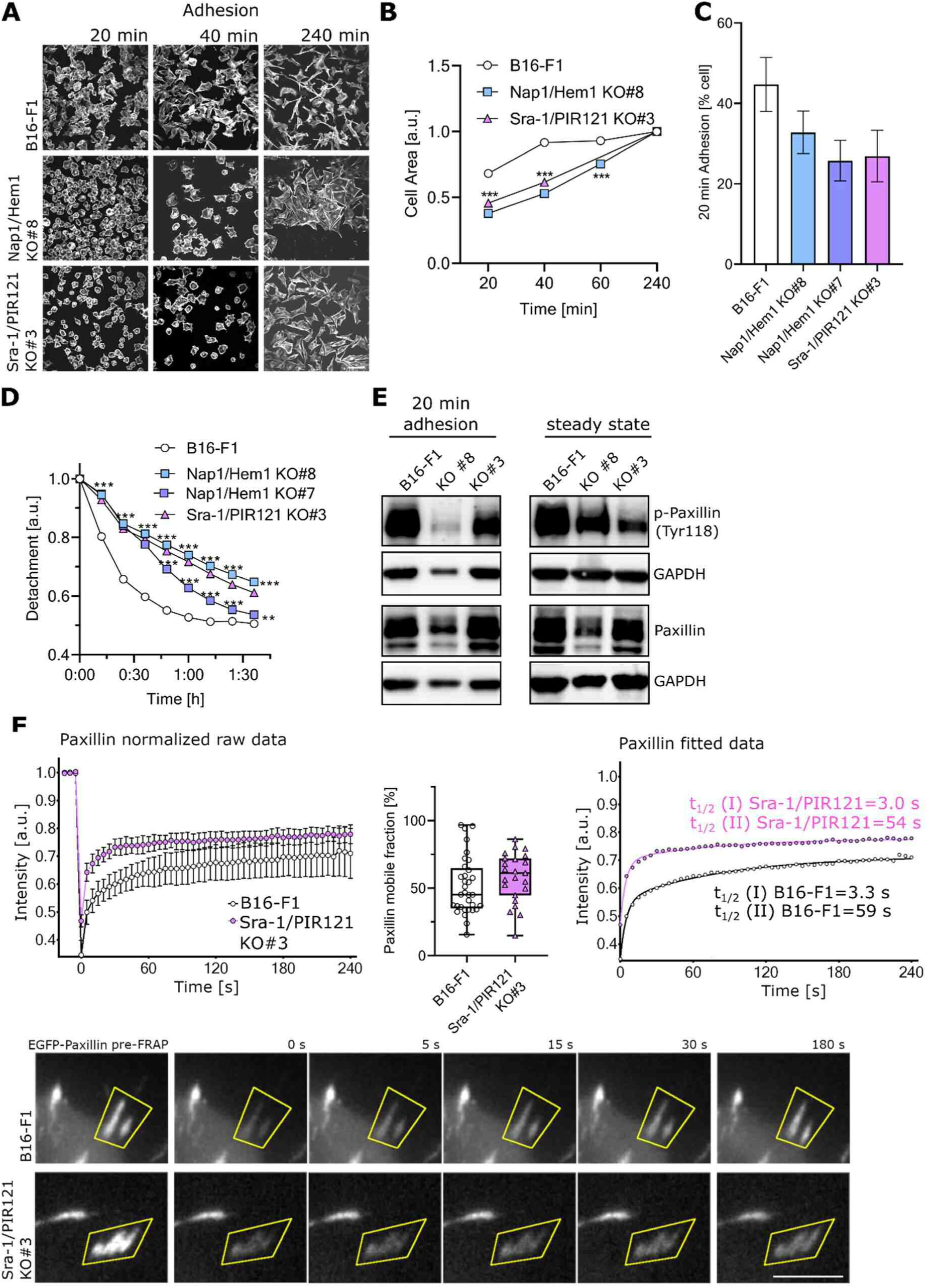
Altered cell adhesion and paxillin dynamics in WRC-deficient B16-F1 cells. **(A)**Immunofluorescent staining of the actin cytoskeleton of B16-F1 cells allowed to adhere on fibronectin. WRC-deficient cells with different genotypes and B16-F1 as controls were analyzed concerning their cell morphologies upon adhesion for different time points as indicated. Bar=50µm **(B)** Quantification of cell area upon spreading on fibronectin for indicated time points. Values are normalized to respective endpoints, confirming delayed spreading in absence of WRC. **(C)** Percentage of B16-F1 cells firmly attached after 20min adhesion to fibronectin (6 replicates) is reduced by approximately 36 % in the absence of functional WRC. **(D)** Detachment assay of B16 cells as measured by decreasing eccentricity (n=2; 6 replicates each), confirming more persistent adhesion upon loss of WRC function. **(E)** Representative Western blots of paxillin and phospho-paxillin expression during steady state and after 20min adhesion on fibronectin. **(F)** FRAP analysis of EGFP-paxillin in individual focal adhesions of B16-F1 genotypes as indicated (WT *versus* Sra-1/PIR121 KO), including determination of the mobile fraction (middle) from normalized raw data (left panel), and the half-time of recovery values as obtained from fitted data (right graph). FRAP behavior of EGFP-paxillin in these experiments best fitted bi-exponential curves according to formula y=Y0+a*(1-exp(-bX))+c(1-exp(-dX), which assumes fluorescence recovery to derive from two populations of molecules, a rapidly and a slowly cyclin population, thus revealing two separable half times of recovery, t ½(I) and t ½(II). The data reveal a modest increase in both the total fraction of cycling paxillin molecules (mobile fraction, middle panel), and the half times of recovery for the two cycling populations by approx. 10% in the absence of WRC (Sra-1/PIR121 KO) in case of the latter (right panel). The bottom panel displays individual bleaching events encased by yellow polygons. Note that EGFP-tagged paxillin was also more difficult to bleach on average in Sra-1/PIR121 KOs, as best illustrated on the raw data panel (top left), indicating rapid exchange with non-bleached molecules already during the short bleaching period (see detailed methods in supplemental data), and adding to the view of an increased mobile fraction seen in the absence of WRC.

### 3. Phagocytosis by macrophages in the absence of WRC

Another hallmark of macrophage function is their ability to remove unwanted material by phagocytosis. This activity, alike cell migration, is driven by a complex interplay of protrusion and adhesion (Dupuy&Caron JCS2008). Phagocytosis, however, comes in different modes depending on the receptors that are engaged by the material to be engulfed. It is well known, for instance, that complement receptor (CR3, consisting of CD11b and CD18, building αMβ2 integrin)-mediated phagocytosis leads to profound Rho-kinase and myosin II activation and is accompanied by Arp2/3 complex-mediated actin assembly (*55, 56*). In contrast, Fc gamma receptor (member of the immunoglobulin superfamily)-mediated phagocytosis was shown to require Rac- and Cdc42 GTPase activation and protrusion to envelop the particle. These results would suggest that Fc gamma receptor-mediated phagocytosis might depend on WRC but not integrins, whereas CR3-triggered phagocytosis prominently involves adhesion, integrin and myosin signaling, but less protrusion (*55*). More recently, Freeman and colleagues found that integrins are also essentially involved in Fc gamma phagocytosis by forming a diffusion barrier around the phagocytic target, coordinating particle engulfment (*57*). We thus first analyzed pathogen-associated (Fig. 4A-D) and FcγR or C3R dependent phagocytic ability (Fig. 4E-G) of WT vs Hem1 KO and WASP KO macrophages. In principle, all macrophage genotypes were capable of internalizing material. Quantifying the phagocytic capability revealed a clear decrease for WASP KO macrophages, in line with published work (*23*). However, phagocytosis defects were most profound in Hem1 KO macrophages, being reduced in phagocytic performance by 50-85% (Fig. 4 A-E & Fig. 4H). The amount of Hem1 KO macrophages that phagocytosed pHrodo green-labeled *E. coli* was decreased by about 85% and 80% in comparison to WT and WASP KO macrophages respectively (Fig. 4A-B). A similar trend was observed using pHrodo green-labeled zymosan particles (Fig. 4C and Movie S4A). We wondered how pathogen phagocytosis may phenotypically appear in the absence of WRC and performed scanning electron microscopy (SEM) of macrophages phagocytizing *E. coli* bacteria. SEM revealed that WT macrophages displayed significant ruffling upon contact with bacteria, leading to their engulfment and finally internalization. In contrast, Hem1 KO macrophages were unable to form canonical phagocytic cups, and particles appeared to sink into the cell instead, with the membrane covering them reminiscent of a membrane pocket (Figs. 4D). We next tested phagocytosis triggered by FcγR by using IgG-opsonized sheep red blood cells (SRBCs). The phagocytic efficiency of Hem1 KO macrophages was again significantly decreased in comparison to both WT and WASP KO macrophages (Fig. 4E, Fig. S4A, Movie S4B). Finally, time lapse microscopy exposed fundamental defects during Hem1 KO macrophage phagocytosis distinct from WASP KO and WT cells. WASP KO macrophages, alike WT, demonstrated increased ruffling as soon as IgG SRBCs contacted the cell, but took generally longer than WT macrophages for fully engulfing them. Uptake of SRBC by WT cells was regularly achieved in less than 2 minutes, frequently engulfing multiple targets at the same time (Fig. S4A and Movie S4B). Hem1 KO macrophages were also observed to engage multiple targets but as opposed to WT, frequently failed to completely phagocytose them. Reminiscent of the phenotype observed by SEM for *E. coli*, the cells did not form a canonical phagocytic cup, but the membrane formed a pocket partially encasing the particle, the closure of which though appeared to be impaired, with the particle slipping out of the pocket on several occasions. Hem1 KO macrophages were thus frequently observed to carry a bulk of IgG-SRBCs on their dorsal surface, unable to complete internalization (Fig. S4A, Movie S4B). Since we had already observed that integrin-based adhesion to fibronectin together with adhesion complex turnover is impeded in Hem1 KO macrophages, we tested whether integrin engagement via CR3 (=Mac-1=integrin αMβ2) binding to complement-opsonized particles (*58*) is affected as well. Indeed, CD11b engagement around IgM/complement-opsonized beads was clearly reduced in Hem1 but not WASP KO macrophages as compared to WT (Fig. 4F and Fig. S4B). Finally, we analyzed the precise appearance of phagocytic cups/pockets in all three genotypes by scanning electron microscopy of beads either being coated with IgG (Fig. S4C) or opsonized with IgM/complement (Fig. 4G; Fig. S4D). Phagocytosis appears to be affected through a combination of reduced envelopment of the beads and reduced adhesion of the plasma membrane to the phagocytic targets. This is in particular evident in the later steps of envelopment, where the WT membrane leaflet in contact with the particle is extremely thin and tightly attached to the bead, whereas the Hem1 KO-membrane does not only display an increased number of filopodia, but also a clearly evident gap between bead and membrane (Fig. 4G), indicating perhaps diminished adhesion to the bead. Our results thus clearly implicate WRC-mediated and Arp2/3 complex-driven actin assembly in either integrin delivery to or integrin activation at sites of protrusion, or both. Quantitative results for the phagocytic performance of the different genotypes in different types of phagocytosis are provided in Fig. 4H.

**Figure 4.**
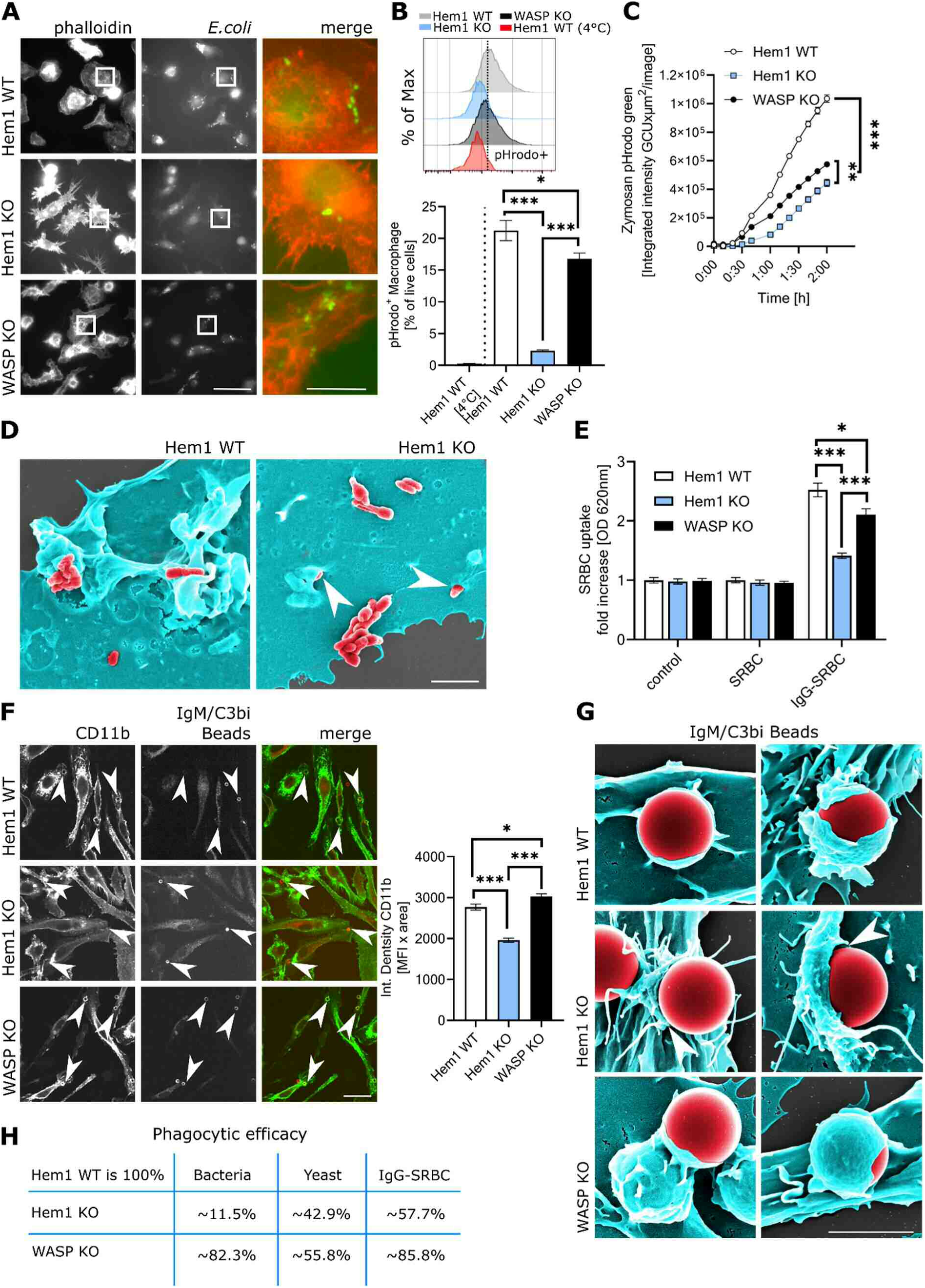
Defective phagocytosis in Hem1 KO macrophages. **(A)**Immunofluorescence images of macrophages challenged with pHrodo™ Green-labelled *E. coli* (green in merge) and stained with phalloidin (red in merge). Bar=50µm and 3µm in the inset. **(B)** Upper panel: Representative histogram overlay of WT (grey), Hem1 KO (blue) and WASP KO (black) macrophages challenged with pHrodo™ Green-labelled *E. coli* (same as in (A)). Bottom curve (red) is the 4°C-control of WT. Lower panel: Quantification of the percentage of pHrodo™ Green positive single, live cells (n=2, 4 replicates). **(C)** Time-resolved phagocytosis by macrophages of pHrodo™ Green-labelled Zymosan bioparticles. Integrated intensity per image is shown over time (n=1; 7 replicates). **(D)** SEM images of macrophages (pseudocolored cyan) phagocytosing pHrodo™ Green labelled E. coli (red). Arrowheads in right image (Hem1 KO) point to membrane pockets formed around bacteria during phagocytosis. Bar=3µm. **(E)** Quantification of phagocytosis of IgG-opsonized sheep red blood cells (SRBCs). Data are normalized to non-treated WT controls (n=3; 8 replicates). **(F)** Left panel: Immunofluorescence of CD11b (green in merge) accumulating at phagocytic cups 2min after challenge with C3bi-opsonized beads (red, d=3µm). Right panel: Quantification of the integrated density of CD11b accumulated around beads. **(G)** SEM images of phagocytosed C3bi-opsonized beads (pseudocolored in red). Arrowheads in the middle panels (Hem1 KO) point to the membrane pockets around beads or gaps between phagocytic membrane and bead (right middle panel). Bar=3µm. **(H)** Direct comparison of quantitative phagocytic performance of macrophages from three genotypes upon challenge with different phagocytic targets.

### 4. Spreading and integrin activation in platelets lacking Hem1

In order to formally establish the intimate connection between WRC–mediated protrusion formation and integrin activation, we turned to another myeloid lineage and analyzed the spreading and adhesion behavior of platelets. Platelets are produced by giant polyploid precursor cells called megakaryocytes (*59*) and play pivotal roles for homeostasis, vascular integrity, as well as inflammatory processes (*60*).

Integrin activation represents the crucial step during platelet activation, enabling their firm adhesion to the injured vessel wall as well as formation of a stable, three-dimensional platelet plug that seals the wound. We therefore investigated integrin function in platelets isolated from WT and Hem1 KO mice at approximately 10-12 weeks of age. In blood smear preparations, red blood cells were confirmed to display a more irregular morphology (*61*). Moreover, significant deviations of the shape of platelets became evident (Fig. 5A). To better understand these changes and in order to relate them to function, we isolated and analyzed WT and Hem1 null platelets. Total counts of platelets from peripheral blood were slightly increased and platelet size was significantly larger in the blood from Hem1 KO mice as compared to the WT (Figure 5B), indicating that Hem 1 is not essential but may play an important role in megakaryocyte differentiation or pro-platelet production and fission *in vivo*. We allowed isolated platelets to spread on glass, which showed that they are in principle capable of spreading, and confirmed their deviant shape. Next, we stained the actin and microtubule cytoskeletons (Fig. S5A), and quantified cells with flat, lamellipodia-like cell edges *versus* spiky shape, analogous to what we had observed with macrophages (compare Fig. 1A). Unexpectedly and in contrast to macrophages, we noticed that Hem1 KO platelets were not completely devoid of lamellipodia, but that approximately one third of the cells displayed either ambiguous protrusions (10%) or even lamellipodia (23%) (Fig. 5C). This result implies that in platelets, but not in macrophages, loss of Hem1 can be bypassed, at least in part, by a yet unknown compensatory mechanism.

**Figure 5.**
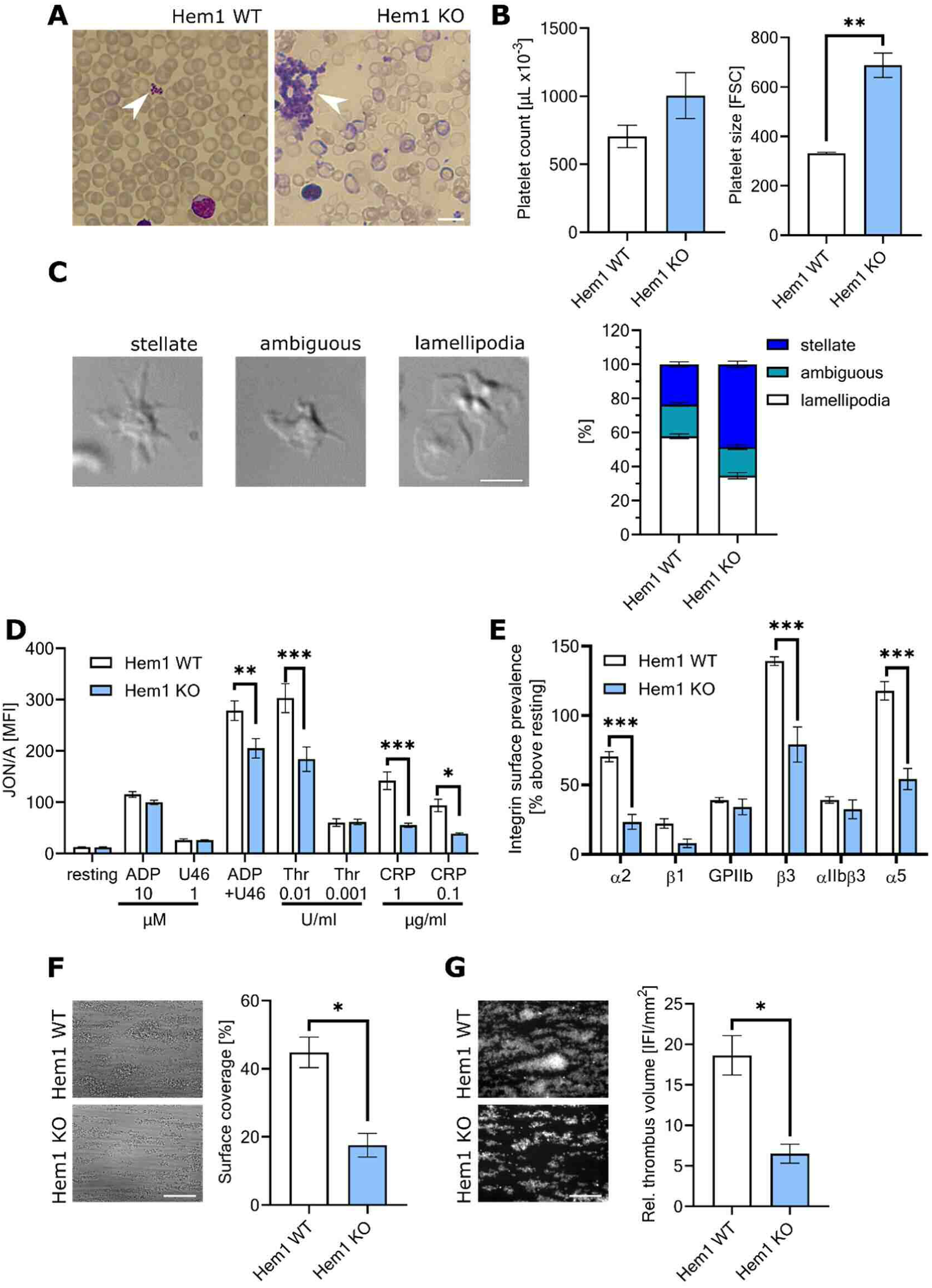
Defective integrin activation in Hem1 KO platelets. **(A)** Giemsa-stained blood smears of Hem1 KO and WT mice confirming affected erythrocytes (*61*) and illustrating abnormal platelet formation (arrow). Bar=5µm. **(B)** Flow cytometric analysis of platelet counts (left) and platelet size (right). **(C)** Left panel: DIC images representing platelet morphology upon stimulation with thrombin and 30min of spreading on fibrinogen. Morphology categorization into “stellate” (Hem1 KO), “ambiguous” (Hem1 KO) and “lamellipodia” (Hem1 WT). Bar=5µm. Right panel: quantification of platelets according to the three morphology categories in WT and Hem1 KO, as indicated (n=3). **(D)** Assessment of platelet activation *in vitro* by flow cytometry of integrin αllbβ3 activation (binding of JON/A-PE antibody) in response to indicated agonists (n=4). ADP: adenosine di-phosphate, Thr: thrombin, U46: U46619, CRP: collagen-related peptide. **(E)** Accumulation of integrins at the platelet plasma membrane upon stimulation with thrombin (0.01 U/ml), as analyzed by flow cytometry and shown as increase of surface prevalence compared to resting (n=4). **(F)** Representative DIC images of whole anti-coagulated blood from WT or Hem1 KO perfused over a collagen-coated surface (0.2 mg/mL) at a shear rate of 1000s-1 (n=5), and quantification of thrombus surface coverage (right panel). **(G)** Representative images of thrombi as in (F), but fluorescently labelled (left). Relative thrombus volume expressed as integrated fluorescence intensity (IFI) per square millimetre (right). Bar=50µm.

Notwithstanding this, flow cytometric analysis of isolated platelets revealed that expression of the most abundant, platelet-specific integrin αIIbβ3 (GPIIbIIIa) was not significantly different between Hem1 null and WT cells (Fig. S5B). However, a slight reduction in the expression of integrin α2 and β1, as well as a more significant reduction of CD9, a tetraspanin that associates with platelet integrins, was apparent. Levels of the von Willebrand Factor (vWF) receptor complex GPIb/IX/V and the collagen receptor GPVI were unchanged or only slightly altered between the two genotypes (Fig. S5B).

Activation with agonists stimulating either G protein-coupled receptors (ADP, thrombin, thromboxane analogue U46619) or GPVI/ITAM signalling leads to inside-out activation of integrins, in particular of the major platelet integrin αIIbβ3, which can be measured by the Jon/A-PE antibody (*62*), preferentially binding the active conformation of αIIbβ3 (Figure 5D). Strikingly, Hem1 null platelets displayed significantly reduced activation of αIIbβ3 in response to GPVI stimulation by collagen-related peptide (CRP). A similar trend to reduced integrin activation was also observed in response to high thrombin concentrations, as well as combined stimulation with ADP and U46619 (Figure 5D). Moreover, surface exposure of the adhesion molecule P-Selectin upon activation, which in resting cells is stored in α-granules, is another hallmark of platelet activation and helps them to adhere to the vessel endothelium. However, degranulation-dependent surface exposure of P-selectin was only mildly affected in response to all stimuli tested (Figure S5C). Finally, agonist-induced upregulation of integrin α2, α5 and β3 prevalence on the platelet surface was found to be also reduced in Hem1 KO platelets (Figure 5E). Thus, the adhesion parameter most conclusively and consistently altered in platelets in the absence of WRC at this stage was αIIbβ3 integrin activation. These results thus indicate a specific defect of integrin-mediated adhesion in the absence of Hem1 in platelets.

To investigate the consequences of defective integrin activation for functionality of Hem1 null platelets under more physiological conditions, we performed an *ex vivo* flow adhesion assay, where whole anti-coagulated blood is perfused over a collagen-coated cover slip at arterial shear rates (Fig. 5 F-G). WT platelets rapidly adhered to the surface and formed stable, three-dimensional aggregates resembling a thrombus. In contrast, thrombi formed by blood of Hem1 null animals displayed both a smaller surface coverage (assessed by thrombus outline, Fig. 5F) and reduced volume (fluorescent intensity, Figure 5G). These findings are in line with our observations that Hem1 KO mice displayed a clear, but not life-threatening bleeding phenotype, as exemplified for instance by increased bleeding during ear-coding. In summary, these results revealed that Hem1 deficiency results not only in expected morphological alterations, but also in prominent integrin activation defects in platelets, significantly compromising their function under static and dynamic conditions.

## DISCUSSION

The work reported here uncovers how WRC- and Arp2/3-driven actin polymerization enable integrin activation to facilitate dynamic adhesion to the substratum or to phagocytic targets and thus support immune cell functions. We also provide new mechanistic insights into the interplay between protrusion and adhesion, concerting effective cell motility.

Within the WASP superfamily of NPFs, WRC is essential for sheet-like protrusions of the plasma membrane such as lamellipodia or ruffles of mesenchymal cells (*63-65*), JAILS of endothelial cells (*66*) or phagocytic cups of immune cells (*67*). WRC exists as various isoform combinations of its five essential subunits (*68*), with Hem1 being the only hematopoietic-specific one (*69, 70*). We had previously analyzed dendritic cells derived from bone marrow of mice genetically lacking Hem1 and consequently WRC function (*7, 9*). In these rapidly migrating leukocytes, actin-driven lamellipodial protrusions facilitated exploration of the environment and directional choices, critical for pathfinding through and invasion of complex matrices. Therefore, DC-migration was reminiscent of bleb-driven locomotion where the cell front is not involved in adhesion and force transduction (*9*). Here, we explored adhesion-dependent motile processes and cell migration, as exerted by macrophages or mesenchymal-like tumor. In such cells, nascent adhesions are frequently born beneath the lamellipodium, mature while the cell moves over them and mediate force coupling between cytoskeleton and substratum, thus facilitating translocation of the cell body (*71*).

We analyzed macrophages and platelets from Hem1 null mice along with somatic cells lacking WRC upon CRISPR/Cas9-mediated deletion of selected WRC components. Loss of WRC leads to a strikingly altered cell morphology with a stellate to spiky cell shape, completely lacking lamellipodia and displaying a surplus of filopodia. Despite this, chemotaxis and migration in 2D environments are only mildly affected. In contrast, migration through increasingly complex matrices/environments is strongly impaired. *In vivo*, this is mirrored by a drastic accumulation of myeloid cells in vessel walls and in the interstitium. We considered that defective matrix degradation might be causative of this phenotype and thus analyzed podosome formation and consecutive matrix degradation, which is not reduced, however, in the absence of WRC. This is in line with earlier studies finding podosome formation to be strictly dependent on WASP-mediated Arp2/3 complex activation (*32, 72*), which is normal in the absence of WRC. Our results thus confirm that WRC is dispensable for podosome and invadopodia formation as well as extracellular matrix degradation. This conclusion supports the previously published observation that suppression of WRC in transformed cells promotes invasive behavior through N-WASP (*73*).

Phagocytosis requires protrusion of the plasma membrane to envelop the target particle, and is well accepted to involve both WASP and WAVE proteins (*13, 23, 67, 74*). Earlier work already established that Arp2/3 complex - albeit not essential - is crucially contributing to phagocytosis (*55, 75*), while the precise differential contributions of WASP and WRC remained elusive. Accordingly, phagocytosis was found here to be detectable even in the absence of Hem1, although the engulfment of different targets such as bacteria, yeast and opsonized red blood cells or beads was significantly delayed or defective to different degrees as compared to WT and WASP KO macrophages. This emphasizes that effective envelopment and timely internalization of target particles by professional phagocytes is strongly supported though not exclusively mediated by WRC-driven protrusion. Different phagocytic targets engage a multitude of corresponding receptors and thus induce different signaling cascades (*76*). Complement-opsonized target particles mediate phagocytosis through complement receptor (CR3), major activation of Rho and subsequent Rho kinase- and myosin II-controlled contraction. In contrast, Fc gamma receptor (FcγR)-induced phagocytosis is mainly accompanied by Rac GTPase signaling and Arp2/3 complex-driven membrane protrusion (*55, 56, 77*). Phagocytosis is in any case a multi-step process involving ligand binding, membrane protrusion, integrin activation and contraction, thus involving both Rac and Rho signaling, albeit to differential degrees and depending on the phagocytic receptor engaged (*40, 78*). This may also explain the observed reduction of phagocytic efficacy of the different targets to variable extents. However, only the three canonical Rac GTPases, aided perhaps by Arf GTPases such as Arf1, are considered capable of activating WRC (*49, 79-82*). Therefore, our results emphasize the clear involvement of the Rac/Arf/WRC/Arp2/3 complex signaling axis in most if not all types of phagocytosis.

It is clear that integrin activation plays a central role during phagocytosis (*40, 83-86*), since most phagocytic receptors embody members of the integrin family. Indeed, Freeman and colleagues demonstrated in an elegant piece of work how integrin activation leads to the formation of a diffusion barrier around the opsonized target, which facilitates coordinated particle engulfment during Fc gamma receptor-initiated phagocytosis (*87*). Interestingly, another recent study using Arp2 knockout cells identified a critical role for Arp2/3 complex-mediated actin assembly in integrin activation, affecting all integrin-dependent processes in macrophages, including migration, adhesion and phagocytosis (*75*). However, the specific participation of single Arp2/3 complex activators could not yet be delineated, since all Arp2/3-mediated processes were simultaneously switched off. Therefore, we analyzed integrin-dependent immune cell functions to unambiguously clarify the specific role of WRC-driven actin assembly in integrin-dependent processes.

The adhesion of Hem1 null macrophages to fibronectin, particularly during cell spreading, was found to be significantly delayed, whereas podosome formation was intact due to the unaltered function of WASP. However, cell detachment took significantly longer in the absence of WRC as compared to WT cells. Adhesions are commonly born as focal complexes within or at the base of lamellipodia and occasionally mature into focal adhesions and finally fibrillary adhesions as the cell moves over them, before they are dissolved at the cell rear. In order to learn whether there is a conserved role for WRC-mediated, Arp2/3 complex-driven lamellipodia formation in integrin activation, we followed several experimental approaches to obtain non-hematopoietic cell lines lacking functional WRC. To this end, we simultaneously removed Hem1 and Nap1 in mouse melanoma cells by CRISPR/Cas9 genome editing, and compared these lines to the previously established lines lacking the Rac-binding WRC-subunits Sra1 and PIR121 (*49*). Strikingly, both types of cells adhered to substrata *and* detached from them with significant delay as compared to WT, further corroborating the assumption of a general role for WRC activity in the regulation of adhesion, as opposed to a Hem1 or Nap1-specific phenotype.

At last, another well-established model for signal-induced spreading and integrin-dependent adhesion with concomitant lamellipodium formation are platelets. This peculiar and important component of blood abundantly expresses the platelet-specific integrin αIIbβ3, which is rapidly activated upon signals such as CRP (*88*), leading to spreading and lamellipodium formation. Assessment of αIIbβ3 integrin activation in platelets derived from Hem1 KO mice uncovered strongly reduced integrin activation despite unchanged αIIbβ3 surface levels (Fig. S5B). Moreover, Hem1-deficient platelets, similar to Hem1 null DCs, macrophages and WRC-deficient B16-F1 cells, displayed severe alterations in cell morphology. However, in contrast to macrophages or dendritic cells, Hem1 null platelets still also showed some pancake-shaped cells and cells with clear lamellipodia, at least at a certain frequency. This was surprising, but we cannot exclude at present that-as opposed to DCs or macrophages - platelets might be capable of expressing low amounts of the mostly non-hematopoietic Hem1 orthologue Nap1, allowing for WRC-driven Arp2/3 complex activation in a fashion indistinguishable from Hem1. Interestingly, platelets conditionally deleted for Cyfip1 but still harboring the intact gene for Cyfip2 display related but less severe phenotypes with aberrant morphology and reduced activatability (*89*), potentially due to residual WRC function. Likewise, when generating mouse melanoma cells CRISPR/Cas9-edited for Nap1, we isolated clones with differential but low levels of compensatory Hem1-expression, giving rise to significant levels of lamellipodia formation (*48*). Thus, Nap1 expression pen etrating into platelets constitutes one potential explanation for the observations made. Nonetheless, platelets lacking Hem1 displayed clearly detectable phenotypes in the majority of cells, concerning cell shape, spreading behavior and integrin activation that perfectly suit our observations in other cell types.

Expression profiling confirmed that cell adhesion is affected upon loss of Hem1 function in macrophages. In addition, protein expression analyses of adhesion components revealed a significant reduction of paxillin phosphorylation on tyrosine 118. In mature adhesions, paxillin is localized close to the plasma membrane, between integrin and the mechanosensitive vinculin/talin complex; it is therefore believed to embody a core component of a molecular clutch between adhesion complexes formed in association with integrin transmembrane receptors and cytoplasmic actin filaments connecting to and likely pulling on these adhesion complexes (Case & Waterman 2015). Paxillin was previously identified as major player in the regulation of adhesion assembly and turnover, becoming enriched and strongly phosphorylated by focal adhesion kinase (FAK) already in early focal complexes (nascent adhesions). In more mature adhesions then, phospho-paxillin is enriched at the distal tip of adhesions and the signal declines in aged adhesions (*47, 53*). Since the speed of both assembly and disassembly of focal adhesions is altered in Hem1 KO cells, along with defects in paxillin phosphorylation, we tested paxillin turnover in adhesions in the absence of WRC. FRAP measurements of EGFP-tagged paxillin turnover in B16-F1 cells gene-deleted for the WRC components Sra-1/PIR121 revealed a modest increase in exchange rates of paxillin in these adhesions. The overall fraction of paxillin capable of turnover in given adhesion sites (mobile fraction) was also increased. Thus, it is tempting to speculate that in the absence of WRC and thus Arp2/3 complex-driven lamellipodial actin networks, paxillin phosphorylation is compromised in the focal complex-initiation zone (normally beneath lamellipodia), leading to reduced retention of paxillin in adhesions. It is known for more than a decade that tyrosine phospho-mimetic or non-phosphorylatable mutants of paxillin affect both assembly and turnover of adhesion sites (*53*). Furthermore, overexpression of a phospho-mimetic mutant (Pax Y13/118E) enhances lamellipodial protrusion and formation of focal complexes and focal adhesions, whereas cells overexpressing a non-phosphorylatable paxillin mutant (Pax Y13/118F) display prominent fibrillary adhesions. Likewise, FAK overexpression triggers focal adhesion disassembly (*53*). All these observations are in line with our results obtained in Hem1 null macrophages and Hem1/Nap1- or Sra-1/Pir121 KO B16-F1 cells. One conclusion may be that lamellipodial actin networks impose spatiotemporal control on the paxillin phosphorylation-cycle and thus its turnover in adhesions. Since FAK was reported to directly bind Arp2/3 complex (*90, 91*), we speculate that in cells devoid of WRC- and Arp2/3 complex-mediated lamellipodia formation, nascent adhesions suffer from a local lack of pre-activated FAK-paxillin complexes. This might in turn contribute to the defective activation of integrins in such adhesions. This hypothesis is supported by our finding that integrin activation is also affected in platelets lacking Hem1.

Finally, the phagocytic defects observed in the absence of Hem1 and thus WRC are also very likely connected, at least in part, to defective integrin activation and thus reduced adhesion to the phagocytic target. It is reasonable to assume that travelling of the protruding plasma membrane over the phagocytic target benefits from stickiness through integrin-based adhesion, believed to be regulated by the same molecular clutch mechanisms as found in focal adhesions (*92*). In agreement with this view, most if not all types of phagocytosis involve one or the other integrin (*40, 57, 76*), rendering phagocytosis a canonical, integrin-dependent process.

We here propose a model, in which both establishment and dissolution of integrin contacts requires the presence of Arp2/3-driven actin structures in their immediate vicinity, facilitating paxillin phosphorylation and inside-out integrin activation. This would work irrespective of how Arp2/3 complex is activated, as exemplified by normal podosome formation in Hem1 KO macrophages. In this scenario, NPFs dictate the location where integrins become activated, at least indirectly (e.g. WASP for podosomes and WRC for focal complexes beneath lamellipodia and during phagocytosis). Hence, the NPF in each case would only indirectly impact on integrin activation by locally generating the accompanying actin structure.

The precise mechanism of integrin activation by Arp2/3 complex-nucleated actin networks, however, remains to be established, although it is attractive to hypothesize the involvement of one of three likely scenarios or any combination of them: (i) the physical interaction of Arp2/3 complex with adhesion components like FAK (*90, 91, 93*), (ii) the mere presence of branched actin filaments, which flow over and directly bind to several adhesion components like talin, zyxin or vinculin (*94*), or (iii) additional factors that become transported to these sites through the respective actin network such as cortactin (*95, 96*), mAbp1 (*97*), filamin (*98, 99*) and/or α-actinin (*100, 101*). These proteins are known to fulfil the criteria by promoting phagocytosis and integrin adhesion, and - in loss of function experiments-share features with the cellular phenotypes obtained upon Hem1 knockout. In any case, future work will have to dissect these possibilities concerning the relative contributions of these players to local integrin activation in addition to their direct, mechanistic function in a given actin structure.

In conclusion, we here establish that WRC also generates lamellipodial actin networks in macrophages that are utilized to reach out for new space during migration and spreading or to envelop particles during phagocytosis. Moreover, we establish that WRC- and Arp2/3 complex-nucleated actin filament networks are crucial for local integrin activation, facilitating adhesion and force coupling of the protruding membrane to the substratum. Loss of the WRC subunit Hem1 leads to a significant delay of cell adhesion and de-adhesion of macrophages, platelets and genome-edited cancer cells. Moreover, a severe defect in phagocytosis in cells lacking Hem1 is associated with the tendency of the phagocytic target to escape the phagocytic cup, indicating lack of (integrin-dependent) stickiness of the enveloping membrane. This has severe consequences for the host, as inefficiently cleared pathogens, apoptotic and necrotic cell bodies are a major source of inflammation and rise of autoantigens, respectively. Indeed, numerous studies have suggested that inefficiently cleared cells can promote the development of autoimmune diseases, such as observed in recently described human patients harboring Hem1 mutations (*102*).

## MATERIALS & METHODS

### Animals

Hem1 KO strain Nckap1l^tm1.2.Sixt^ (MGI ID: 6197558)(*9*) and WASP KO strain WAS^tm1Kas^ (MGI ID: 3028420)(*23*) were bred on C57BL/6 (Jackson Laboratories) background at the animal facility of the Helmholtz Centre for Infection Research, Braunschweig, Germany. Mice were kept under SPF conditions in individually ventilated cages and fed *ad libitum*. Mice were sacrificed via CO2 inhalation to harvest cells and organs. Mice were handled in accordance with the national animal welfare law.

### Macrophage differentiation and tissue culture procedures

Bone marrow-differentiated macrophages were isolated from the femurs of Hem1 WT, Hem1 KO and WASP KO mice. Bone marrow-derived cells were differentiated for a minimum of 7 days and further cultivated for a maximum of 21 days in macrophage medium (RPMI, 10%FBS (v/v) and 20% (v/v) conditioned L929 medium), as described (*103*). Peritoneal exudate cells were isolated by flushing the peritoneum of mice with 5ml lavage buffer. Peritoneal cells were centrifuged (1200rpm, 4°C, 5min), resuspended in macrophage medium and left to settle on culture plates. Cells were washed to remove suspension cells 1h after seeding, and peritoneal macrophages were grown at 7.5% CO2 and 37°C. Cultivation did not exceed 2 days.

### Maintenance and CRISPR/Cas9-mediated genome editing of B16-F1 cells

B16-F1 cells lacking both NCKAP1L (Hem1) and NCKAP1 (Nap1) genes were generated using CRISPR/Cas9 technology as described (*104*), but starting with a cell line already lacking functional Nap1 (*48*). Hem1 knockout was subsequently achieved on the Nap1 null background with CRISPR-guide sequence 5’-CTCACGATCCTGAATGACCG-3. Cell clones were screened for the absence of both gene products via Western Blot and confirmed by genomic sequencing. B16-F1 cells (ATCC CRL-6323) and Nap1/Hem1 or Sra1/PIR121 double knockout clones (*49*) derived thereof were cultured in DMEM supplemented with 10% FBS (v/v) and 2mM L-glutamine.

### Platelet preparation

Washed platelets were prepared as described (*60*). Briefly, mice were anesthetized using isoflurane and bled into 300µl TBS buffer containing 20U/ml heparin. The blood was centrifuged (twice, 300g, 6min) to obtain platelet-rich plasma (PRP). PRP was supplemented with 0.02U/ml apyrase and 0.1µg/ml PGI2 and platelets were pelleted (800g, 5min) and washed twice with Tyrode-HEPES buffer supplemented with 0.02U/ml apyrase and 0.1µg/ml PGI2. Platelets were allowed to rest 30min prior to experiments.

### Cremaster preparation, staining and spinning disk confocal microscopy

Cremaster preparation and staining were mainly performed as described (*37*). Briefly, the scrotum of sacrificed male mice was sterilized and carefully opened to expose the testis. The cremaster was pre-fixed *in situ* with 4% PFA for 15min at RT and then carefully excised from the tissue. Next, the muscle was expanded, mounted in position and fixed again with 4% PFA for 1h at RT. Permeabilization occurred with 0.5% TX 100 (1h, RT). Incubation with primary antibodies took place overnight. Secondary antibody incubation occurred for 1h at RT. Next, samples were washed in 0.05% TX 100 and mounted in Prolong™ Gold Antifade reagent. Spinning disk confocal microscopy was performed with a Perkin Elmer Ultra View spinning disk microscope, and images were taken in Z stack intervals of 0.5µm with a 60x Plan Apochromat immersion objective lens (1.4 NA). Interval counts were adjusted for the individual thickness of the sample. The software Volocity (Quorum Technologies, Canada) was used for image acquisition and analysis.

### Cell biological experiments and microscopy

Scanning EM was performed as described in (*105*). Fluorescence microscopy of fixed and stained cells, video Microscopy & FRAP experiments were performed as described in (*106*). Platelet Assays were essentially done as described in (*107*). Flow cytometry was performed with a FACS Aria (Becton Dickinson) following standard procedures. Images were adjusted for brightness and contrast with ImageJ software (NIH). Detailed procedures for each experiment are also provided in the supplemental material and all reagents used are listed in Table S2.

### Data presentation and statistical analysis

Data presentation and statistical evaluation was performed in GraphPad Prism 8 software (*P≤0.05, **P≤0.0l, ***P≤0.000l). If not otherwise indicated, data are presented as means and standard error of means (mean+SEM). Error bars were not depicted if the error bar was shorter than symbol size. For simple comparison between 3 groups, ordinary one way ANOVA with Tukey’s multiple comparison test was employed. For grouped analyses, two-way ANOVA with Tukey’s multiple comparison was employed. Non-parametric Mann-Whitney rank sum test was employed in case of comparing only 2 sample groups with each other. Lack of statistical significance was not indicated.

## SUPPLEMENTAL MATERIAL

Figure S1. Unaltered differentiation and activation of Hem1 KO macrophages *in vitro*.

Figure S2. Differential regulation of focal adhesion dynamics in Hem1 null macrophages.

Figure S3. Altered adhesion component regulation in the absence of WRC

Figure S4. FcγR- and CR3-dependent phagocytosis

Figure S5. Cytoskeleton staining and adhesion molecule surface patterns of Hem1 KO platelets

Movie S1A Random migration of macrophages

Movie S1B 3D reconstruction of cremaster preparation Movie S2 Detachments Assay of Macrophages

Movie S3A Detachment assay of B16-F1 cells Movie S3B FRAP of EGFP-tagged paxillin Movie S3C FRAP of EGFP-tagged VASP

Movie S4A Phagocytosis of pHrhodo labelled zymosan particles Movie S4B phagocytosis of IgG opsonized SRBCs

Table 1: List of genes regulated in Hem1 KO versus WT macrophages Supplemental Methods

Table S2. List of all materials used in this study

## ACKNOWLEDGMENTS

We are grateful to Silvia Prettin, Ina Schleicher and Petra Hagendorff for expert technical assistance. We thank the HZI animal facility and David Dettbarn for excellence in animal keeping and breeding. We thank Werner Tegge for peptides used in immunization and Giorgio Scita for antibodies. This work was supported, in part, by the Deutsche Forschungsgemeinschaft (DFG), Priority Programm SPP1150 (to TS, KR, JF and MSi), and by DFG-grant GRK2223/1 (to KR). TS was also supported by the Helmholtz Society HGF impulse fund W2/W3-066. MSc is supported by the Mexican Council for Science and Technology (CONACyT, 284292), Fund SEP-Cinvestav (108), and the Royal Society, UK (Newton Advanced Fellowship, NAF/R1/180017).

## Author contributions

SS, TS, CK and KR conceptualized and drafted figures for the manuscript. SS, TS, KR, AS, MSi, MSc, RG and BN contributed to experimental design and interpretation of results. SS, HD, CK, DG, SD, AG, MR, MM, FK, AS, JF conducted experimental procedures and analyzed data. SS, TS, KR and IP wrote the original draft. All authors participated in editing and revising the manuscript text and figures.

## Competing interests

The authors declare no competing interests.

## Data and materials availability

All data supporting the findings of this study are available within the article. Hem1 KO mice are available upon reasonable request to TS and dependent on a mutually signed material transfer agreement.

## REFERENCES

1. J. D. Humphries, A. Byron, M. J. Humphries, Integrin ligands at a glance. J Cell Sci 119, 3901–3903 (2006).

2. S. T. Pals, D. J. de Gorter, M. Spaargaren, Lymphoma dissemination: the other face of lymphocyte homing. Blood 110, 3102–3111 (2007).

3. G. Giannone et al., Lamellipodial actin mechanically links myosin activity with adhesion-site formation. Cell 128, 561–575 (2007).

4. D. A. Lauffenburger, A. F. Horwitz, Cell migration: a physically integrated molecular process. Cell 84, 359–369 (1996).

5. T. J. Mitchison, L. P. Cramer, Actin-based cell motility and cell locomotion. Cell 84, 371–379 (1996).

6. T. Lammermann et al., Rapid leukocyte migration by integrin-independent flowing and squeezing. Nature 453, 51–55 (2008).

7. H. R. Thiam et al., Perinuclear Arp2/3-driven actin polymerization enables nuclear deformation to facilitate cell migration through complex environments. Nat Commun 7, 10997 (2016).

8. R. Alon, M. L. Dustin, Force as a facilitator of integrin conformational changes during leukocyte arrest on blood vessels and antigen-presenting cells. Immunity 26, 17–27 (2007).

9. A. Leithner et al., Diversified actin protrusions promote environmental exploration but are dispensable for locomotion of leukocytes. Nat Cell Biol 18, 1253–1259 (2016).

10. T. D. Pollard, G. G. Borisy, Cellular motility driven by assembly and disassembly of actin filaments. Cell 112, 453–465 (2003).

11. J. V. Small, T. Stradal, E. Vignal, K. Rottner, The lamellipodium: where motility begins. Trends Cell Biol 12, 112–120 (2002).

12. P. Kunda, G. Craig, V. Dominguez, B. Baum, Abi, Sra1, and Kette control the stability and localization of SCAR/WAVE to regulate the formation of actin-based protrusions. Curr Biol 13, 1867–1875 (2003).

13. H. Park et al., A point mutation in the murine Hem1 gene reveals an essential role for Hematopoietic protein 1 in lymphopoiesis and innate immunity. The Journal of experimental medicine 205, 2899–2913 (2008).

14. A. Steffen et al., Filopodia formation in the absence of functional WAVE- and Arp2/3-complexes. Molecular biology of the cell 17, 2581–2591 (2006).

15. M. Sixt, M. Bauer, T. Lammermann, R. Fassler, Beta1 integrins: zip codes and signaling relay for blood cells. Current opinion in cell biology 18, 482–490 (2006).

16. M. L. Dustin, T. A. Springer, T-cell receptor cross-linking transiently stimulates adhesiveness through LFA-1. Nature 341, 619–624 (1989).

17. J. C. Nolz et al., The WAVE2 complex regulates actin cytoskeletal reorganization and CRAC-mediated calcium entry during T cell activation. Curr Biol 16, 24–34 (2006).

18. P. A. Zipfel et al., Role for the Abi/wave protein complex in T cell receptor-mediated proliferation and cytoskeletal remodeling. Curr Biol 16, 35–46 (2006).

19. M. H. Pauker et al., WASp family verprolin-homologous protein-2 (WAVE2) and Wiskott-Aldrich syndrome protein (WASp) engage in distinct downstream signaling interactions at the T cell antigen receptor site. J Biol Chem 289, 34503–34519 (2014).

20. J. C. Nolz et al., The WAVE2 complex regulates T cell receptor signaling to integrins via Abl- and CrkL-C3G-mediated activation of Rap1. The Journal of cell biology 182, 1231–1244 (2008).

21. J. C. Nolz et al., WAVE2 regulates high-affinity integrin binding by recruiting vinculin and talin to the immunological synapse. Molecular and cellular biology 27, 5986–6000 (2007).

22. E. Rivers, A. J. Thrasher, Wiskott-Aldrich syndrome protein: Emerging mechanisms in immunity. Eur J Immunol 47, 1857–1866 (2017).

23. J. Zhang et al., Antigen receptor-induced activation and cytoskeletal rearrangement are impaired in Wiskott-Aldrich syndrome protein-deficient lymphocytes. The Journal of experimental medicine 190, 1329–1342 (1999).

24. S. Benesch et al., N-WASP deficiency impairs EGF internalization and actin assembly at clathrin-coated pits. J Cell Sci 118, 3103–3115 (2005).

25. M. Innocenti et al., Phosphoinositide 3-kinase activates Rac by entering in a complex with Eps8, Abi1, and Sos-1. The Journal of cell biology 160, 17–23 (2003).

26. M. Innocenti et al., Abi1 regulates the activity of N-WASP and WAVE in distinct actin-based processes. Nat Cell Biol 7, 969–976 (2005).

27. S. Gordon, Alternative activation of macrophages. Nat Rev Immunol 3, 23–35 (2003).

28. S. R. Paludan, Interleukin-4 and interferon-gamma: the quintessence of a mutual antagonistic relationship. Scand J Immunol 48, 459–468 (1998).

29. K. A. Jablonski et al., Novel Markers to Delineate Murine M1 and M2 Macrophages. PLoS One 10, e0145342 (2015).

30. D. Ishihara, A. Dovas, H. Park, B. M. Isaac, D. Cox, The chemotactic defect in wiskott-Aldrich syndrome macrophages is due to the reduced persistence of directional protrusions. PLoS ONE 7, e30033 (2012).

31. G. E. Jones, D. Zicha, G. A. Dunn, M. Blundell, A. Thrasher, Restoration of podosomes and chemotaxis in Wiskott-Aldrich syndrome macrophages following induced expression of WASp. Int J Biochem Cell Biol 34, 806–815 (2002).

32. S. Linder, D. Nelson, M. Weiss, M. Aepfelbacher, Wiskott-Aldrich syndrome protein regulates podosomes in primary human macrophages. Proc Natl Acad Sci U S A 96, 9648–9653 (1999).

33. A. J. Thrasher, S. Burns, R. Lorenzi, G. E. Jones, The Wiskott-Aldrich syndrome: disordered actin dynamics in haematopoietic cells. Immunol Rev 178, 118–128 (2000).

34. S. Tsuboi, A complex of Wiskott-Aldrich syndrome protein with mammalian verprolins plays an important role in monocyte chemotaxis. J Immunol 176, 6576–6585 (2006).

35. D. Zicha et al., Chemotaxis of macrophages is abolished in the Wiskott-Aldrich syndrome. Br J Haematol 101, 659–665 (1998).

36. B. M. Isaac et al., N-WASP has the ability to compensate for the loss of WASP in macrophage podosome formation and chemotaxis. Exp Cell Res 316, 3406–3416 (2010).

37. J. Song et al., Endothelial Basement Membrane Laminin 511 Contributes to Endothelial Junctional Tightness and Thereby Inhibits Leukocyte Transmigration. Cell Rep 18, 1256–1269 (2017).

38. D. S. Harburger, D. A. Calderwood, Integrin signalling at a glance. J Cell Sci 122, 159–163 (2009).

39. I. Kourtzelis, I. Mitroulis, J. von Renesse, G. Hajishengallis, T. Chavakis, From leukocyte recruitment to resolution of inflammation: the cardinal role of integrins. J Leukoc Biol 102, 677–683 (2017).

40. A. G. Dupuy, E. Caron, Integrin-dependent phagocytosis: spreading from microadhesion to new concepts. J Cell Sci 121, 1773–1783 (2008).

41. S. Linder, M. Aepfelbacher, Podosomes: adhesion hot-spots of invasive cells. Trends Cell Biol 13, 376–385 (2003).

42. R. Buccione, J. D. Orth, M. A. McNiven, Foot and mouth: podosomes, invadopodia and circular dorsal ruffles. Nature reviews 5, 647–657 (2004).

43. S. Linder, P. Kopp, Podosomes at a glance. J Cell Sci 118, 2079–2082 (2005).

44. A. M. Lopez-Colome, I. Lee-Rivera, R. Benavides-Hidalgo, E. Lopez, Paxillin: a crossroad in pathological cell migration. J Hematol Oncol 10, 50 (2017).

45. K. Nakamura et al., Tyrosine phosphorylation of paxillin alpha is involved in temporospatial regulation of paxillin-containing focal adhesion formation and F-actin organization in motile cells. J Biol Chem 275, 27155–27164 (2000).

46. C. E. Turner, K. M. Pietras, D. S. Taylor, C. J. Molloy, Angiotensin II stimulation of rapid paxillin tyrosine phosphorylation correlates with the formation of focal adhesions in rat aortic smooth muscle cells. J Cell Sci 108 (Pt 1), 333–342 (1995).

47. R. Zaidel-Bar, C. Ballestrem, Z. Kam, B. Geiger, Early molecular events in the assembly of matrix adhesions at the leading edge of migrating cells. J Cell Sci 116, 4605–4613 (2003).

48. S. Dolati et al., On the relation between filament density, force generation, and protrusion rate in mesenchymal cell motility. Molecular biology of the cell 29, 2674–2686 (2018).

49. M. Schaks et al., Distinct Interaction Sites of Rac GTPase with WAVE Regulatory Complex Have Non-redundant Functions in Vivo. Curr Biol 28, 3674–3684 e3676 (2018).

50. C. Ballestrem, B. Hinz, B. A. Imhof, B. Wehrle-Haller, Marching at the front and dragging behind: differential alphaVbeta3-integrin turnover regulates focal adhesion behavior. The Journal of cell biology 155, 1319–1332 (2001).

51. C. Ballestrem, B. Wehrle-Haller, B. A. Imhof, Actin dynamics in living mammalian cells. J Cell Sci 111 (Pt 12), 1649–1658 (1998).

52. I. Chandrasekar et al., Vinculin acts as a sensor in lipid regulation of adhesion-site turnover. J Cell Sci 118, 1461–1472 (2005).

53. R. Zaidel-Bar, R. Milo, Z. Kam, B. Geiger, A paxillin tyrosine phosphorylation switch regulates the assembly and form of cell-matrix adhesions. J Cell Sci 120, 137–148 (2007).

54. B. Stutchbury, P. Atherton, R. Tsang, D. Y. Wang, C. Ballestrem, Distinct focal adhesion protein modules control different aspects of mechanotransduction. J Cell Sci 130, 1612–1624 (2017).

55. R. C. May, E. Caron, A. Hall, L. M. Machesky, Involvement of the Arp2/3 complex in phagocytosis mediated by FcgammaR or CR3. Nat Cell Biol 2, 246–248 (2000).

56. I. M. Olazabal et al., Rho-kinase and myosin-II control phagocytic cup formation during CR, but not FcgammaR, phagocytosis. Curr Biol 12, 1413–1418 (2002).

57. S. A. Freeman et al., Integrins Form an Expanding Diffusional Barrier that Coordinates Phagocytosis. Cell 164, 128–140 (2016).

58. V. Jaumouille, A. X. Cartagena-Rivera, C. M. Waterman, Coupling of beta2 integrins to actin by a mechanosensitive molecular clutch drives complement receptor-mediated phagocytosis. Nat Cell Biol 21, 1357–1369 (2019).

59. K. R. Machlus, J. E. Italiano, The incredible journey: From megakaryocyte development to platelet formation. The Journal of cell biology 201, 785–796 (2013).

60. B. Nieswandt, I. Pleines, M. Bender, Platelet adhesion and activation mechanisms in arterial thrombosis and ischaemic stroke. J Thromb Haemost 9 Suppl 1, 92–104 (2011).

61. M. M. Chan et al., Hematopoietic protein-1 regulates the actin membrane skeleton and membrane stability in murine erythrocytes. PLoS One 8, e54902 (2013).

62. W. Bergmeier et al., Flow cytometric detection of activated mouse integrin alphaIIbbeta3 with a novel monoclonal antibody. Cytometry 48, 80–86 (2002).

63. M. Innocenti et al., Abi1 is essential for the formation and activation of a WAVE2 signalling complex. Nat Cell Biol 6, 319–327 (2004).

64. L. M. Machesky, R. H. Insall, Scar1 and the related Wiskott-Aldrich syndrome protein, WASP, regulate the actin cytoskeleton through the Arp2/3 complex. Curr Biol 8, 1347–1356 (1998).

65. A. Steffen et al., Sra-1 and Nap1 link Rac to actin assembly driving lamellipodia formation. EMBO J 23, 749–759 (2004).

66. A. Abu Taha, M. Taha, J. Seebach, H. J. Schnittler, ARP2/3-mediated junction-associated lamellipodia control VE-cadherin-based cell junction dynamics and maintain monolayer integrity. Molecular biology of the cell 25, 245–256 (2014).

67. R. Lorenzi, P. M. Brickell, D. R. Katz, C. Kinnon, A. J. Thrasher, Wiskott-Aldrich syndrome protein is necessary for efficient IgG-mediated phagocytosis. Blood 95, 2943–2946 (2000).

68. O. Alekhina, E. Burstein, D. D. Billadeau, Cellular functions of WASP family proteins at a glance. J Cell Sci 130, 2235–2241 (2017).

69. R. Hromas, S. Collins, W. Raskind, L. Deaven, K. Kaushansky, Hem-1, a potential membrane protein, with expression restricted to blood cells. Biochimica et Biophysica Acta (BBA) - Gene Structure and Expression 1090, 241–244 (1991).

70. O. D. Weiner et al., Hem-1 complexes are essential for Rac activation, actin polymerization, and myosin regulation during neutrophil chemotaxis. PLoS Biol 4, e38 (2006).

71. L. B. Case, C. M. Waterman, Integration of actin dynamics and cell adhesion by a three-dimensional, mechanosensitive molecular clutch. Nat Cell Biol 17, 955–963 (2015).

72. S. Burns, A. J. Thrasher, M. P. Blundell, L. Machesky, G. E. Jones, Configuration of human dendritic cell cytoskeleton by Rho GTPases, the WAS protein, and differentiation. Blood 98, 1142–1149 (2001).

73. H. Tang et al., Loss of Scar/WAVE complex promotes N-WASP- and FAK-dependent invasion. Curr Biol 23, 107–117 (2013).

74. A. M. Pearson et al., Identification of cytoskeletal regulatory proteins required for efficient phagocytosis in Drosophila. Microbes Infect 5, 815–824 (2003).

75. J. D. Rotty et al., Arp2/3 Complex Is Required for Macrophage Integrin Functions but Is Dispensable for FcR Phagocytosis and In Vivo Motility. Dev Cell 42, 498–513 e496 (2017).

76. S. A. Freeman, S. Grinstein, Phagocytosis: receptors, signal integration, and the cytoskeleton. Immunol Rev 262, 193–215 (2014).

77. E. Caron, A. Hall, Identification of two distinct mechanisms of phagocytosis controlled by different Rho GTPases. Science 282, 1717–1721 (1998).

78. H. Park, D. Cox, Cdc42 regulates Fc gamma receptor-mediated phagocytosis through the activation and phosphorylation of Wiskott-Aldrich syndrome protein (WASP} and neural-WASP. Molecular biology of the cell 20, 4500–4508 (2009).

79. B. Chen et al., Rac1 GTPase activates the WAVE regulatory complex through two distinct binding sites. Elife 6, (2017).

80. K. Kobayashi et al., p140Sra-1 (specifically Rac1-associated protein) is a novel specific target for Rac1 small GTPase. J Biol Chem 273, 291–295 (1998).

81. V. Koronakis et al., WAVE regulatory complex activation by cooperating GTPases Arf and Rac1. Proc Natl Acad Sci U S A 108, 14449–14454 (2011).

82. V. Singh, A. C. Davidson, P. J. Hume, D. Humphreys, V. Koronakis, Arf GTPase interplay with Rho GTPases in regulation of the actin cytoskeleton. Small GTPases 10, 411–418 (2019).

83. D. I. Beller, T. A. Springer, R. D. Schreiber, Anti-Mac-1 selectively inhibits the mouse and human type three complement receptor. The Journal of experimental medicine 156, 1000–1009 (1982).

84. C. Cougoule, A. Wiedemann, J. Lim, E. Caron, Phagocytosis, an alternative model system for the study of cell adhesion. Semin Cell Dev Biol 15, 679–689 (2004).

85. R. Hanayama et al., Identification of a factor that links apoptotic cells to phagocytes. Nature 417, 182–187 (2002).

86. J. Savill, I. Dransfield, N. Hogg, C. Haslett, Vitronectin receptor-mediated phagocytosis of cells undergoing apoptosis. Nature 343, 170–173 (1990).

87. S. A. Freeman et al., Applied stretch initiates directional invasion through the action of Rap1 GTPase as a tension sensor. J Cell Sci 130, 152–163 (2017).

88. S. Pasco, J. C. Monboisse, N. Kieffer, The alpha 3(lV}185-206 peptide from noncollagenous domain 1 of type IV collagen interacts with a novel binding site on the beta 3 subunit of integrin alpha Vbeta 3 and stimulates focal adhesion kinase and phosphatidylinositol 3-kinase phosphorylation. J Biol Chem 275, 32999–33007 (2000).

89. Y. Schurr et al., Platelet lamellipodium formation is not required for thrombus formation and stability. Blood 134, 2318–2329 (2019).

90. B. Serrels et al., Focal adhesion kinase controls actin assembly via a FERM-mediated interaction with the Arp2/3 complex. Nat Cell Biol 9, 1046–1056 (2007).

91. V. Swaminathan, R. S. Fischer, C. M. Waterman, The FAK-Arp2/3 interaction promotes leading edge advance and haptosensing by coupling nascent adhesions to lamellipodia actin. Molecular biology of the cell 27, 1085–1100 (2016).

92. S. A. Freeman, S. Grinstein, Phagocytosis: Mechanosensing, Traction Forces, and a Molecular Clutch. Curr Biol 30, R24–R26 (2020).

93. K. A. DeMali, C. A. Barlow, K. Burridge, Recruitment of the Arp2/3 complex to vinculin: coupling membrane protrusion to matrix adhesion. The Journal of cell biology 159, 881–891 (2002).

94. D. L. Huang, N. A. Bax, C. D. Buckley, W. I. Weis, A. R. Dunn, Vinculin forms a directionally asymmetric catch bond with F-actin. Science 357, 703–706 (2017).

95. S. McFarlane, C. McFarlane, N. Montgomery, A. Hill, D. J. Waugh, CD44-mediated activation of alpha5beta1-integrin, cortactin and paxillin signaling underpins adhesion of basal-like breast cancer cells to endothelium and fibronectin-enriched matrices. Oncotarget 6, 36762–36773 (2015).

96. F. Agerer et al., Cellular invasion by Staphylococcus aureus reveals a functional link between focal adhesion kinase and cortactin in integrin-mediated internalisation. J Cell Sci 118, 2189–2200 (2005).

97. J. Schymeinsky et al., A fundamental role of mAbp1 in neutrophils: impact on beta(2) integrin-mediated phagocytosis and adhesion in vivo. Blood 114, 4209–4220 (2009).

98. C. D. Lynch et al., Filamin depletion blocks endoplasmic spreading and destabilizes force-bearing adhesions. Molecular biology of the cell 22, 1263–1273 (2011).

99. E. Berrou et al., Gain-of-Function Mutation in Filamin A Potentiates Platelet Integrin alphaIIbbeta3 Activation. Arterioscler Thromb Vasc Biol 37, 1087–1097 (2017).

100. S. Tadokoro et al., A potential role for alpha-actinin in inside-out alphaIIbbeta3 signaling. Blood 117, 250–258 (2011).

101. F. Jahan et al., Phosphorylation of the alpha-chain in the integrin LFA-1 enables beta2-chain phosphorylation and alpha-actinin binding required for cell adhesion. J Biol Chem 293, 12318–12330 (2018).

102. W. A. Comrie et al., Genetic immunodeficiency and autoimmune disease reveal distinct roles of Hem1 in the WAVE2 and mTORC2 complexes. bioRxiv 692004, (2019).

103. V. Trouplin et al., Bone marrow-derived macrophage production. J Vis Exp, e50966 (2013).

104. F. Kage et al., FMNL formins boost lamellipodial force generation. Nat Commun 8, 14832 (2017).

105. M. G. Coppolino et al., Evidence for a molecular complex consisting of Fyb/SLAP, SLP-76, Nck, VASP and WASP that links the actin cytoskeleton to Fcgamma receptor signalling during phagocytosis. J Cell Sci 114, 4307–4318 (2001).

106. A. Steffen et al., Rac function is crucial for cell migration but is not required for spreading and focal adhesion formation. J Cell Sci 126, 4572–4588 (2013).

107. I. Pleines et al., Multiple alterations of platelet functions dominated by increased secretion in mice lacking Cdc42 in platelets. Blood 115, 3364–3373 (2010).

108. B. Nieswandt, W. Bergmeier, K. Rackebrandt, J. E. Gessner, H. Zirngibl, Identification of critical antigen-specific mechanisms in the development of immune thrombocytopenic purpura in mice. Blood 96, 2520–2527 (2000).

109. D. M. Mosser, X. Zhang, Measuring opsonic phagocytosis via Fcgamma receptors and complement receptors on macrophages. Curr Protoc Immunol Chapter 14, Unit 14 27 (2011).

110. K. Rottner, M. Krause, M. Gimona, J. V. Small, J. Wehland, Zyxin is not colocalized with vasodilator-stimulated phosphoprotein (VASP} at lamellipodial tips and exhibits different dynamics to vinculin, paxillin, and VASP in focal adhesions. Mol Biol Cell 12, 3103–3113 (2001).

111. U. D. Carl et al., Aromatic and basic residues within the EVH1 domain of VASP specify its interaction with proline-rich ligands. Curr Biol 9, 715–718 (1999).

