## Supplemental Material and Figures for "Loss of Hem1 disrupts macrophage function and impacts on migration, phagocytosis and integrin-mediated adhesion"

**This file includes:**

Supplemental methods

Table S2. Key resource table

Fig. S1. Unaltered differentiation and activation of Hem1 KO macrophages *in vitro*

Fig. S2. Differential regulation of focal adhesion dynamics in Hem1 null macrophages

Fig. S3. Altered adhesion component regulation in the absence of WRC

Fig. S4. FCγR- and CR3-dependent phagocytosis

Fig. S5. Cytoskeleton staining and adhesion molecule surface patterns of Hem1 KO platelets

Legends for supplemental movies S1A-S4B

References (108-111)

### Supplemental Material

#### Supplemental Methods

##### *Western blotting*

Cells were counted using a Luna II system (BioCat, Germany), seeded and treated as indicated in respective experiment. For whole cell lysates, cells were washed with ice-cold PBS, taken up in 2x SDS sample buffer and boiled (5min, 95°C). Western blotting of equal amounts of protein lysates was carried out using standard techniques. Primary and secondary antibodies are listed in Table S2. Chemiluminiscence signals were obtained upon incubation with Lumi-Light Western Blotting Substrate in combination with the INTAS ECL Chemocam Imager (Intas Science Imaging, Germany). Exposure times were adjusted for each antibody. Signals were quantified by densitometry with ImageJ. Background staining was reduced using the ImageJ plugin Rolling Ball Background Subtraction. Average intensities of bands of interest were normalized to respective loading control.

##### *Immunostaining*

Macrophages and B16-F1 cells were seeded onto fibronectin-coated glass coverslips, treated as indicated and fixed in pre-warmed 4% paraformaldehyde (PFA) in PBS for 20min. Next, cells were blocked and permeabilized in 5% horse serum, 1% BSA and 0.1% saponin in PBS for 1h at RT. Cells were subsequently stained with primary antibodies as indicated (Table S2) for 1h at RT. All antibody solutions contained 5% horse serum, 1% BSA and 0.1% saponin. The procedure was repeated with respective secondary antibodies (Table 1). Afterwards, coverslips were washed in PBS and mounted in Prolong™ Diamond Antifade Mountant with DAPI to additionally stain cell nuclei. For visualization of the actin cytoskeleton, either Alexa488-conjugated phalloidin, ATTO594-conjugated phalloidin or ATTO647-conjugated phalloidin was employed.

##### *Platelet spreading assay*

Coverslips were coated with 200µg/ml fibrinogen overnight at 4°C. Platelets were stimulated with 0.01U/ml thrombin and allowed to spread. Platelets were fixed with 4% PFA in PBS for 30min and visualized with a Zeiss Axiovert 200M inverted microscope (100x/ 1.4 oil objective with differential interference contrast (DIC)). Digital images were recorded using a CoolSNAP-EZ camera (Visitron Systems GmbH, Puchheim, Germany). For analysis of the distributions of filamentous actin (F-actin) and α-tubulin by immunofluorescence confocal microscopy, platelets were fixed in PHEM buffer supplemented with 4% PFA and 1% IGEPAL® CA-630, and stained as indicated. Samples were mounted in Fluoroshield (Abcam), and images acquired with a Leica TCS SP5 confocal microscope (Leica Microsystems, Germany).

***Platelet adhesion under flow conditions***

Rectangular coverslips (15x60mm) were coated over night with 200µg/ml collagen at 37°C, washed with PBS and blocked in 1% BSA in PBS for 1h at 37°C. Blood was anticoagulated using heparin and further diluted using Tyrode-HEPES buffer supplemented with 2mM $CaCl_2$, incubated with 0.2µg/ml Dylight-488-conjugated anti-GPIX derivate, generated and modified as described (*108*), at 37°C for 5min. Transparent flow chambers with slit depth of 50µm, equipped with the coated coverslips, were connected to the blood-filled syringe. Perfusion was performed at shear stress equivalent to a wall shear rate of $1000s^{-1}$. Blood was perfused for 4min over the collagen-coated surface and washed with Tyrode-HEPES buffer for 4min. Time-lapse phase contrast and fluorescence images were recorded with a Zeiss Axiovert 200 inverted microscope (40x/0.60NA objective) equipped with a CoolSNAP-EZ camera (Visitron, Germany), and analyzed offline using MetaVue software (Molecular Devices, USA).

***Macrophage and B16-F1 spreading assay***

Cells were detached with 1mM EDTA, counted and seeded at $1x10^5$cells on fibronectin-coated (25µg/ml) coverslips (d=12mm). Centrifugation (500rpm, 1min) ensured simultaneous contact of cells with coverslips. Cells were fixed and stained with phalloidin after indicated time points. Images were taken with a Nikon Ti2 eclipse microscope (40x oil objective). Individual areas were measured by thresholding of phalloidin staining using NIS Elements software (Nikon, Japan). Areas were normalized to the average area measured after 4h spreading time for cells from each genotype.

***Macrophage and B16-F1 adhesion assay***

Cells were detached with 1mM EDTA in PBS, resuspended in respective medium and stained with 0.1µM of the viability marker Calcein-AM (37°C, 30min). Cell were washed in PBS, counted, resuspended in RPMI or DMEM and centrifuged to the bottom of 96-well plates (500rpm, 1min) to allow for synchronized adhesion, while being incubated in an IncuCyte S3® system (Sartorius, Germany). The total amount of living cells was measured 20min after adhesion by imaging Calcein-AM-positive cells with the IncuCyte® Analysis software. Next, cells were washed and attached Calcein-AM-positive cells measured again. Adhesion efficacy was quantified by calculating the percentage of attached cells after washing relative to the amount of total cells. All analyses were done using the IncuCyte S3® software.

***Detachment assay***

Macrophages and B16-F1 cells were seeded ($5x10^4$ cells) onto 96-well plates one day before the experiment. On the day of experiment, cells were stained with 0.1µM Calcein-AM and subsequently washed with PBS. 100µl 1mM EDTA in PBS was added to 96-well plates, and cells were immediately transferred into the IncuCyte S3® system (Sartorius). Detachment of cells was followed every 10min for 2h. Detachment was identified by rounding of live cells as defined by their eccentricity (eccentricity of a circle is defined as zero). Change of eccentricity over time was normalized to the initial value of cells from respective genotype. All analyses were performed with the IncuCyte S3® software (see movie S3A).

***Time-lapse microscopy for phagocytosis and 2D/3D chemotaxis assays***

Time lapse microscopy was performed with a Nikon Eclipse Ti E inverted microscope system (Nikon, Japan) with a stage incubation chamber (Okolab, Italy) set to 37°C, 7.5% $CO_2$, 90% humidity. Images were acquired with 10x CFI60 objective and NIS Elements software every 10min for 20h. At least 20 randomly selected cells were tracked per experiment and analyzed with the TrackMate and Chemotaxis Tool plugins of ImageJ software. 2D-directed migration experiments were performed with 100ng/ml C5α as chemoattractant (R&D Systems, USA) using the µ Slide Chemotaxis chamber (Ibidi, Germany) according to manufacturer's instructions. For 3D directed migration assays towards 100ng/ml C5α, macrophages were seeded into a 2mg/ml collagen I matrix. Briefly, for 200µl of 2mg/ml collagen I matrix, 80µl of 5mg/ml rat tail collagen I was mixed with 20µl 10x RPMI on ice. The mixture was neutralized with 5µl 1N NaOH, and 95µl $3x10^6$ cells/ml macrophage suspension was added for a final volume of 200µl. The mixture was carefully dispensed in a µ Slide 4 well Ph+ glass bottom dish (Ibidi) in such a way that half of the chamber was covered in the collagen matrix/cell mixture. The collagen matrix was polymerized over the next 30min in the incubator. Next, the other half of the chamber was filled with a similarly prepared collagen matrix that received the chemoattractant instead of the cell suspension. The collagen matrix was polymerized for 30min and then overlaid with RPMI 1640 to prevent dehydration. Observation area was the border between collagen/cell matrix and collagen/chemoattractant. At least 3 observation points per sample were imaged every 10min for 20h.

##### *Fluorescence recovery after photobleaching (FRAP)*

Time-lapse imaging was performed using an inverted Axio Observer (Zeiss) with a 100x/1.4 NA PLAN-Apochromat oil immersion objective. Bleaching of EGFP-paxillin and EGFP-VASP was conducted using the 2D-VisiFRAP Realtime Scanner controlling a 405 nm laser (Visitron Systems GmbH, Puchheim, Germany) at 90-100 mW power output. Acquired images were adjusted for contrast and brightness using MetaMorph software (Molecular Devices). Data analysis and processing were carried out essentially as described in (*105*).

##### *ECM degradation assay*

2mg/ml FITC-coupled gelatin was prepared by labelling porcine gelatin with FITC according to manufacturer's instructions. Glass coverslips were coated with 0.5µg/ml poly-L-lysine (PLL) for 20min at RT and crosslinked with 0.5% glutaraldehyde in PBS for 15min at RT. 0.2% FITC-coupled gelatin in 2% sucrose in PBS was added and incubated in the dark for 10min, and subsequently quenched with freshly prepared 5mg/ml sodium borohydride in PBS. Afterwards, coverslips were washed thrice with PBS and incubated in macrophage medium at 37°C before adding peritoneal macrophages. The cells were incubated for 20h. Degradation resulted in dark areas within the FITC-coupled gelatin surface beneath the macrophages. Subsequently, the cells were fixed and stained as indicated. Images were taken by conventional epifluorescence microscopy. Analysis was performed using Metamorph software (Molecular Devices, USA). Degradation capability was calculated as degraded area per cell area.

##### *Flow cytometry of macrophages and platelets*

Single cell suspensions were blocked with CD16/32 containing LIVE/DEAD™ Fixable Blue Dead Cell Stain for 30min at RT. Cells were stained first for extracellular antigens on ice in PBS containing 2% FBS and 1mM EDTA. After staining, cells were washed and fixed with 2% PFA for 20min at RT. In case of intracellular antibody staining, cells were permeabilized with 0.5% Saponin for 15min at RT. Intracellular antibody staining occurred for 30min at RT in the dark. Flow cytometry was performed using the BD LSRII SORP system (BD Bioscience, USA), and data analysis done with FlowJo software. For all antibodies, single stainings were performed and compensation was carried out. FMOs (fluorescence minus one) were performed for all intracellular antibodies. For platelet analysis, 50µl of blood were withdrawn from mice under isoflurane anesthesia, washed with Tyrode-HEPES buffer, diluted (1:20) in Tyrode-HEPES buffer containing 2mM $CaCl_2$ and activated with the indicated agonists and concentrations (15 minutes at 37°C). Activation of αIIbβ3-integrin and P-Selectin were determined using fluorophore-conjugated antibodies. Integrin recruitment to the platelet surface upon thrombin activation was determined as the percentage of fluorophore-conjugated antibodies bound to activated platelets as compared to resting platelets. For glycoprotein expression, heparinized whole blood was diluted (1:20) in Tyrodes-HEPES buffer and incubated with saturating amounts of fluorophore-conjugated antibodies for 15min at room temperature. After the indicated time points, the reactions were stopped by the addition of 500µL of Tyrode-HEPES buffer containing 2mM $CaCl_2$. Analyses were performed on a FACSCalibur (BD Biosciences, USA) flow cytometer. Antibodies were generated and modified in our laboratory as described previously (*108*).

***Macrophage polarization assay***

Macrophage polarization potential towards M1 or M2 state was assessed by flow cytometric analysis. Briefly, macrophages were seeded in triplicates onto 96-well plates and allowed to adhere overnight. Subsequently, macrophages were washed and stimulated for 24h with either 20ng/ml IL-4, 50ng/ml C5α, 100U/ml IFNγ and 10ng/ml LPS or 100U/ml IFNγ and 100ng/ml LPS together. Cells were subsequently washed and detached with ice cold PBS containing 1mM EDTA; staining procedure was performed as described above and analyzed with a LSR II SORP (BD Biosciences, USA). In particular, extracellular staining of CD38 and intracellular staining of Egr2 was used to denote M1 or M2 polarization, respectively. M0 macrophages were classified as expressing neither marker and considered un-polarized.

***Phagocytosis assays***

Bone marrow-derived macrophages were either seeded on glass coverslips for microscopic analysis or on 96-well plates for the IncuCyteS3 and absorbance analysis or 6-well plates for flow cytometric analysis, stimulated with 50ng/ml IFNγ and 10ng/ml LPS overnight, and then challenged with either pHrodo™ Green-labelled *E. coli* or pHrodo™ Green Zymosan Bioparticles (Thermo Fisher Scientific, USA).

E. coli phagocytosis: pHrodo™ Green-labeled *E. coli* were prepared according to the manufacturer's instructions. Pre-stimulated macrophages were challenged with pHrodo™ Green -labeled, heat-inactivated *E. coli* for 20min. Flow cytometry samples were detached with ice cold PBS containing 1mM EDTA, stained for live/dead and analyzed with a LSR II SORP (BD Biosciences). As negative control, WT macrophages were incubated at 4°C and challenged with pHrodo™ Green-labelled *E. coli*. Yeast phagocytosis: Macrophages were challenged 1:10 with pHrodo™ Green Zymosan Bioparticles. The IncuCyte S3® system was used for image acquisition (see movie S4_phrodo zymosan). The total integrated intensity of green fluorescent objects per image at indicated time points was analyzed with IncuCyte S3® software. FcγR-dependent phagocytosis: FcγR-dependent phagocytosis assay using sheep red blood cells (SRBC) or polysterene latex beads (d=3µm) was adapted from D. Mosser and X. Zhang (*109*). Briefly, SRBCs or beads were washed with PBS and opsonized with anti SRBC IgG for 1h at RT. Macrophages were challenged in a ratio of 1:10 and incubated for 20min. Non-phagocytosed SRBCs were lysed by treatment with ACK-lysis buffer for no more than 10min. Macrophages were then lysed completely by treatment with 100µl/well 0.1% SDS in PBS for 10min. 50µl of the macrophage lysate was mixed with 50µl of freshly prepared 10mg/ml 2,7-diaminofluorene in acetic acid, supplemented with 3% hydrogen peroxide. Absorbance at 620nm was measured and normalized to untreated WT control. CR3-dependent phagocytosis and quantification of CD11b: For C3R-dependent phagocytosis, polystyrene beads (d=3µm) were washed and opsonized with rabbit IgM for 1h at RT, washed and additionally incubated with mouse serum for 1h at 37°C. C3R-dependent phagocytosis was performed in serum-free RPMI 1640 to minimize non-specific targeting. Beads were centrifuged on cells at 500rpm for 30s to ensure simultaneous contact, incubated for 2min and fixed immediately. Cells were not permeabilized, and stained for the surface integrin receptor CD11b. IgM/C3bi-opsonized beads were visualized by staining for IgM. Images were taken with a Nikon Ti2 eclipse (Nikon, Japan) and spinning disk technology from Yokogawa (Yokogawa Electric, Japan) in Z-stack intervals of 0.3µm with a 60x immersion objective. Analysis of the integrated density (MFIxarea) of CD11b in direct contact with opsonized beads was done with NIS Elements software (Fig. S4B). In detail, a binary layer was created over the beads and the zone of influence extended by 0.33µm. A second binary layer was created for the CD11b staining. Only areas with a direct overlay of both binary layers (beads and CD11b) were analyzed in each Z-plane.

***Scanning electron microscopy***

For scanning electron microscopy, cells were grown on fibronectin-coated cover slips, treated with heat-inactivated *E. coli* DH5α or beads, and fixed after 20 minutes (*E. coli*) or 2 minutes (beads) with 5% formaldehyde and 2% glutaraldehyde in HEPES buffer for at least 2h. Samples were washed in TE buffer and dehydrated on ice with a graded series of acetone/ethanol (10%, 30%, 50%, 70%, and 90%) for 10min each, followed by two steps in 100% acetone/ethanol at RT. Acetone was used for *E. coli* samples, while ethanol was used for the beads, since the overall shape of beads was affected by acetone. After dehydration, samples were subjected to critical-point drying with liquid $CO_2$ (CPD 30, Bal-Tec, Liechtenstein). Dried samples were fixed onto aluminum stubs with plastic conductive carbon cement (Plano, Germany), and covered with a gold palladium film by sputter coating (SCD 500 Bal-Tec, Liechtenstein). Examinations were performed with a field emission scanning electron microscope Zeiss Merlin (Zeiss, Germany), using the Everhart Thornley HESE2 detector and the inlens SE detector in a 25:75 ratio and an acceleration voltage of 5 kV.

***DNA microarray***

Quality and integrity of total RNA was controlled on Agilent Technologies 2100 Bioanalyzer (Agilent Technologies; Waldbronn, Germany). RNA was extracted using Qiagen RNAeasy according to manufacturer's manual. 2-10ng of total RNA were used for biotin labelling according GeneChip® Pico Kit (Affymetrix). 5.5µg of biotinylated cDNA were fragmented and placed in a hybridization cocktail containing four biotinylated hybridization controls (BioB, BioC, BioD, and Cre) as recommended by the manufacturer. Samples were hybridized to an identical lot of Affymetrix Clariom™ S (400 Format) for 17 hours at 45°C. Hybridization was done for 16h at conditions recommended by the manufacturer. Clariom™ S chips were washed and stained in the Affymetrix Fluidics Station 450. GeneChips were scanned using the Affymetrix GCS 3000. Image analysis was done by Affymetrix® GeneChip® Command Console® Software (AGCC) and Affymetrix® Expression Console™ Software. Raw data obtained after image analysis were further analysed by R/BioConducter packages "oligo" and "Biobase". Raw signal intensities of each probeset (gene feature) were summarized by median polish method. Summarized probeset data was log2 transformed followed by RMA normalization procedure. Finally, the obtained data set was annotated by NetAffx (Affymetrix). Normalized data sets were filtered for informative genes (showing at least expression values > log2(10) in more than two samples). Datasets were tested across all groups (ANOVA) or pairwise using linear models to assess differential expression in context of the multifactorial designed experiment. For statistical analysis and assessing differential expression the R/BioConductor package "limma" was used, utilizing an empirical Bayes method to moderate the standard errors of the estimated log-fold changes. Functional analysis was performed by R package "clusterProfiler".

***Generation of polyclonal antibodies***

Polyclonal antibodies against Hem1 were obtained by immunizing female white New Zealand rabbits with a mixture of three Keyhole limpet hemocyanin (KLH)-conjugated Hem1 peptides (peptide sequences: C-AIANLKADNSSPEEEYKVACL, C-SSLKGYSKRVADIKESKEHAITNSGQFHC and C-MLESCFPYVLLRNAYREVSRAFYLN) at a 1:1:1 ratio together with Freund complete adjuvants (Sigma) followed by five boosts with Freund incomplete adjuvants (Sigma). Specificity of the antibodies was determined in solid phase assays and in immunoblots using wild-type and mutant cells.

| Reagent/Resource | Identifier | Application | Source or Composition |
|---|---|---|---|
| **Cell culture media** | | | |
| DMEM [+]4.5g/L D-Glucose, L-Glutamine [-] Pyruvate | 41965-039 | | Gibco |
| HBSS | L2035 | | Biochrom |
| L929-conditioned medium | | 20% (v/v) | Self-made (supernatant of L929 cells cultured in DMEM+10% FBS (v/v)) |
| Mouse serum | S2160 | | Biowest |
| PBS | 10010-056 | | Gibco |
| RPMI 1640 [+] L-Glutamine | 21875-034 | | Gibco |
| **Cell lines** | | | |
| B16-F1 | ATCC CRL-6323 | | ATCC |
| Nap1 Hem1 KO clone #7 | | | Frieda Kage |
| Nap1 Hem1 KO clone #8 | | | Frieda Kage |
| L929 | ATCC® CCL1TM | | ATCC |
| Sra-1/PIR121 KO clone #3 | | | Mathias Schaks (*49*) |
| **Buffers, chemicals and supplements** | | | |

| | | | |
|---|---|---|---|
| 2x SDS sample buffer | | | 0.5M Tris, 10% SDS (w/v), 20% glycerol, 100mM β-mercaptoethanol, 0.02% bromophenol blue (w/v) |
| ACK-lysis buffer | | Erythrocyte lysing buffer solution | 150mM $NH_4Cl$, 10mM $KHCO_3$, 0.1mM $Na_2EDTA$, pH 7.4 |
| Adenosine 5′-diphosphate (ADP) | A2754 | 10µM | Sigma-Aldrich |
| Apyrase | A6132 | 0.02U/ml | Sigma-Aldrich |
| BSA | A6588 | 1% (w/v) | PanReac AppliChem |
| Calcein-AM | C1430 | 0.1µM | Thermo Fisher Scientific |
| Collagen I | CI48 | 2mg/ml | Playtypus Technologies |
| Collagen-related peptide (CRP) | | 1 or 0.1 µg/ml | gift from Paul Bray (Baylor College, USA) |
| EDTA solution | 03690 | 1mM | Sigma-Aldrich |
| FBS, Qualified (heat-inactivated) | 10270-106 | 10% (v/v) | Gibco |
| Fibrinogen | F4483 | 200µg/ml | Sigma-Aldrich |
| Fibronectin(pure) | 11051407001 | 25µg/ml | Sigma-Aldrich |
| FITC (isomer I) | F7250 | | Sigma-Aldrich |
| Fluoroshield™ | F6182 | | Sigma-Aldrich |
| Glutaraldehyde | G6257 | 0.5% (v/v) | Sigma-Aldrich |
| Heparin | | 20U/ml | Ratiopharm |
| HEPES buffer | | | 0.1M HEPES, 0.09M sucrose, 10mM $CaCl_2$, 10mM MgCl2, pH 6.9 |
| Horm® collagen | Horm® | 200µg/ml | Takeda |
| Horse serum | | 5% (v/v) | Cytogen |
| IGEPAL® CA-630 | I8896 | 1% (v/v) | Sigma-Aldirch |
| Immersion oil type F | MXA22168:30cc | | Nikon |
| Latex beads, polystyrene, 3.0 µm mean particle size | LB30 | 1:10 | Sigma-Aldrich |
| Lavage buffer | | Scavenging of peritoneal exudate cells | 2% FBS, 1mM EDTA in PBS |
| L-Glutamine | 25030-081 | 2mM | Gibco |
| LIVE/DEAD™ Fixable Blue Dead Cell Stain | L23105 | | Thermo Fisher Scientific |
| LPS (from *E. coli*; O127:B8) | L3129 | 10ng/ml | Sigma-Aldrich |

| | | | |
|---|---|---|---|
| Lumi-Light Western Blotting Substrate | 12015200001 | | Roche |
| PFA | A3813 | 4% IF<br>2% FC | PanReac AppliChem |
| PGI2 | P6188 | 0.1µg/ml | Sigma-Aldrich |
| PHEM buffer | | | 60mM PIPES, 25mM HEPES, 10mM EGTA, 2mM $MgCl_2$, pH 6.9 |
| pHrodo™ Green-labeled *E. coli* | P35381 | | Thermo Fisher Scientific |
| pHrodo™ Green Zymosan Bioparticles | P35365 | | Thermo Fisher Scientific |
| Pig skin gelatin | 48720 | | Sigma-Aldrich |
| Poly-L-lysine (PLL) | P8920 | | Sigma-Aldrich |
| Prolong™ Diamond Antifade Mountant with DAPI | S36973 | | Thermo Fisher Scientific |
| ProLong™ Gold Antifade Mountant | P36930 | | Thermo Fisher Scientific |
| Recombinant Mouse Complement Component C5a Protein | 2150-C5 | 100ng/ml | R&D Systems |
| Recombinant mouse IFNγ | 12343534 | 100U/ml | ImmunoTools |
| Recombinant mouse IL-4 | 12340045 | 20ng/ml | ImmunoTools |
| Saponin | 47036 | 0.1% (w/v) | Sigma-Aldrich |
| Sheep red blood cells, packed 10% | IC10-0210-15ml | 1:10 | Innovative Research |
| TBS buffer | | | 20mM Tris-HCl, 137mM NaCl in $H_2O$ |
| TE buffer | | | 20mM TRIS, 1mM EDTA, pH6.9 |
| Thrombin | 10602400001 | 0.01U/ml or 0.01U/ml | Roche |
| Tyrode-HEPES buffer | | | 134mM NaCl, 0.34mM Na2HPO4, 2.9mM KCl, 12mM NaHCO3, 5mM HEPES, 5mM glucose, 0.35%BSA, pH 7.4 |
| U46619 | BML-PG023 | 1µM | Enzo Life Sciences |

| | | | |
|---|---|---|---|
| (Prostaglandin $H_2$/thromboxane $A_2$ receptor agonist) | | | |
| **Plastics** | | | |
| µ Slide Chemotaxis | 80306 | | Ibidi |
| µ-Slide 4 Well Ph+ Glass Bottom | 80447 | | Ibidi |
| Cell counting slides | L12001-LG | for Luna II system | BioCat |
| FITC | F7250 | | Sigma-Aldrich |
| Gelatin from pig skin | G-13187 | | Invitrogen |
| **Software** | | | |
| FlowJo software | | Flow cytometry analysis | FlowJo |
| ImageJ | | Image procession | NIH |
| IncuCyte® Analysis Software | | Image acquisition/analysis | Sartorius |
| Metamorph | | Image analysis | Molecular Devices |
| MetaVue | | Image acquisition/analysis | Molecular Devices |
| NIS Elements AR | | Image acquisition/analysis | Nikon |
| GraphPad Prism 8 | | Data presentation/ statistical analysis | GraphPad Software |
| Volocity | | Image acquisition/ analysis | Quorum Technologies |
| **Phalloidin conjugates** | | | |
| Alexa488 phalloidin | A12379 | | Thermo Fisher Scientific |
| ATTO594 phalloidin | AD594-8 | | ATTO-TEC |
| ATTO647 phalloidin | AD647-8 | | ATTO-TEC |
| **Recombinant DNA** | | | |
| pEGFP-Paxillin | pEGFP-N1_Paxillin | | Anika Steffen (*105, 110*) |
| pEGFP-VASP | pEGFP-C2_VASP | | Anika Steffen (*105, 111*) |

| **Antibodies** | | | | |
|---|---|---|---|---|
| **Target** | **Conjugate** | **Identifier or clone** | **Application** | **Source** |
| Abi-1 | / | W8.3 | 1:100 IF | Giorgio Scita (*63*) |
| Abi-1 | / | E3B1 | 1:2000 WB | Giorgio Scita (*63*) |
| CD11b | FITC | M1/70 | 1:400 FC<br>1:200 IF | BioLegend |

| | | | | |
|---|---|---|---|---|
| CD16/32 | / | 93 | 1:500 FC | BioLegend |
| CD31 | / | ab28364 | 1:400 IF | Abcam |
| CD38 | PE | 90 | 1:2000 FC | BioLegend |
| Egr2 | APC | Erongr2 | 1:100 | eBioscience |
| F4/80 | PE/Dazzle | BM8 | 1:200 FC | BioLegend |
| F4/80 | Brilliant Violet 711 | BM8 | 1:100 FC | BioLegend |
| GAPDH | / | CB1001 | 1:10000 WB | Calbiochem |
| Goat | Peroxidase | 705-035-003 | 1:10000 WB | Dianova |
| Hem1 | / | 149 | 1:10000 WB | Jan Faix (MHH) |
| HSPC300 | / | SC-390459 | 1:1000 WB | Santa Cruz |
| Integrin αIIβ3 (high affinity confirmation) | PE | Jon/A | FC | Emfret |
| Integrin β1 | FITC | HMβ1-1 | FC | BioLegend |
| Mouse | Peroxidase | 115-035-068 | 1:10000 WB | Dianova |
| Mouse | Alexa Fluor 488 | A11029 | 1:400 IF | Invitrogen |
| Mouse | Alexa Fluor 597 | A11032 | 1:400 IF | Invitrogen |
| Nap1 | / | Nap1#4953 | 1:500 | Anika Steffen (*65*) |
| Paxillin | / | 349 | 1:1000 WB | BD Biosciences |
| Phospho-Paxillin (Y118) | / | ab4833 | 1:2000 WB | Abcam |
| P-selectin | FITC | WUG 1.9 | FC | Nieswandt lab (*108*) |
| Rabbit | Peroxidase | 111-035-045 | 1:10000 WB | Dianova |
| Rabbit | Alexa Fluor 488 | A11034 | 1:400 IF | Invitrogen |
| Rabbit | Alexa Fluor 594 | A11037 | 1:400 IF | Invitrogen |
| Rat | Alexa Fluor 488 | A11006 | 1:400 IF | Invitrogen |
| Sra-1 | / | 30A4H4 | 1:1000 WB | Anika Steffen (*65*) |
| SRBC (IgG fraction serum) | / | 55806 | 1:1600 | MP Biomedicals |
| SRBC (IgM fraction serum) | / | CL9000-M | 1:400 | Cedarlane |
| Tubulin | Alexa Fluor 488 | B-5-1-2 | 1:300 IF | Invitrogen |
| Vinculin | / | hVin-1 | 1:5000 WB | Sigma |

| | | | | |
|---|---|---|---|---|
| | | | 1:250 IF | |
| WASP | / | AF3070 | 1:1000 WB | R&D Systems |
| WAVE2 (C14) | / | SC-10394 | 1:1000 WB | Santa Cruz |

**Table S2: Key resource table. IF-immunofluorescence, WB-Western Blot, FC-flow cytometry**

#### Supplemental figures

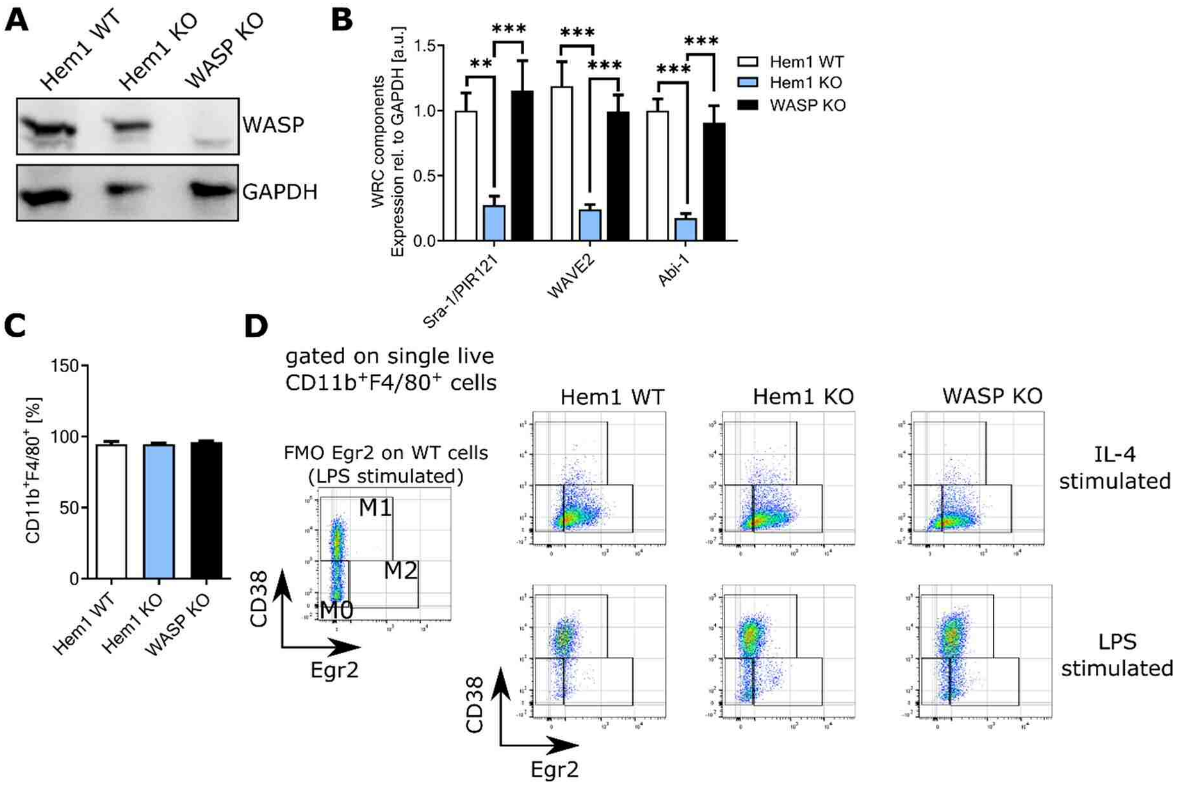


**Figure S1. Unaltered differentiation and activation of Hem1 KO macrophages *in vitro***

**(A)** Western blot showing WASP expression in the indicated macrophage genotypes. GAPDH serves as loading control. Note that loss of Hem1 does not affect WASP expression. **(B)** Quantification of relative expression of WRC components in macrophages in the indicated genotypes (n=5). **(C)** Percentage of macrophages from all bone marrow-derived cells of the three genotypes identified by CD11b$^{+}$F4/80$^{+}$ after 7 days of differentiation *in vitro*. **(D)** Gating strategy to delineate M0/M1/2 polarization states. Left panel shows FMO gating control for Egr2. Right panel exemplifies flow cytometric dot blots for IL-4 and LPS stimulation (n=3).

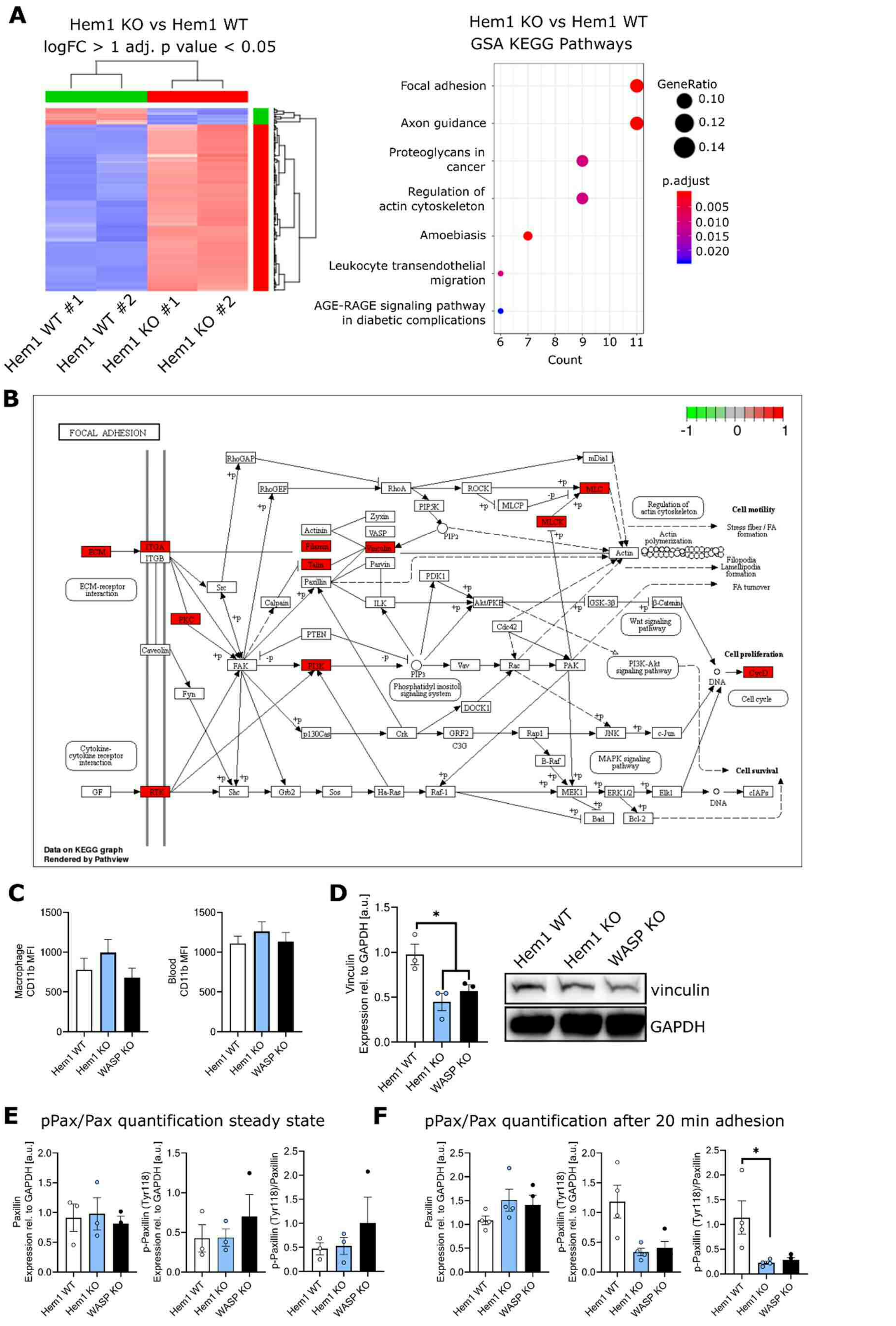


**Figure S2. Differential regulation of focal adhesion dynamics in Hem1 null macrophages**

**(A)** Left panel: Heatmap of RNA expression profile of Hem1 KO versus WT bone marrow derived-macrophages from 2 individual mice each, and right panel: top 7 KEGG (Kyoto Encyclopedia of Genes and Genomes) pathways of differentially expressed genes. **(B)** KEGG pathway for focal adhesion-associated genes with those marked in colour, which are most regulated in Hem1 null *versus* WT macrophages. **(C)** Mean fluorescent intensity of CD11b of bone marrow-derived macrophages (right) (n=3) and of blood cells (n=3), as measured by flow cytometry. **(D)** Western blot and quantification of vinculin expression in macrophages. **(E)** Quantification of paxillin and phospho-paxillin levels during steady state (n=3). **(F)** Quantification of paxillin and phospho-paxillin levels after 20min adhesion to fibronectin (n=4).

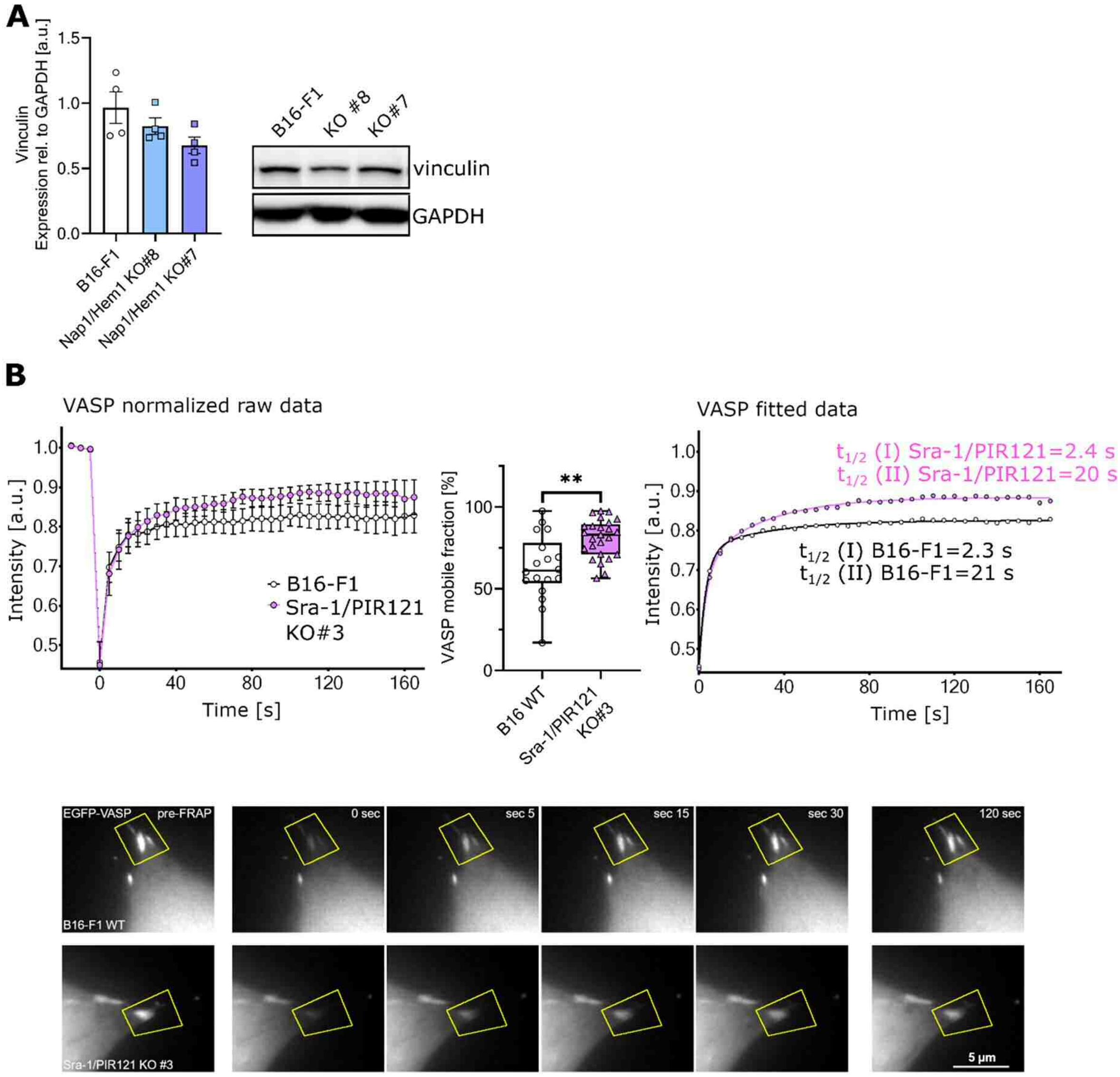


**Figure S3. Altered adhesion component regulation in the absence of WRC**

**(A)** Vinculin expression is reduced in B16-F1 cells following removal of functional WRC (independent of the subunit inactivated, Hem1/Nap1 or Sra-1/PIR121) as exemplified by quantitative Western Blotting (n=4). **(B)** FRAP analysis of individual focal adhesions from GFP-VASP transfected B16-F1 cells (WT and Sra-1/PIR121 KO), with determination of the mobile fraction in the normalized raw data (left panel) and of the two calculated half-time-of-recovery values from the fitted model (right graph). The mobile fraction is defined as the fraction of fluorescence maximally recovered relative to the average intensity measured at bleaching time point 0. Fitted data corresponds to bi-exponential curves according to formula: y=Y0+a*(1-exp(-bX))+c(1-exp(-dX). Note that the mobile fraction of VASP is increased by approx. 22% in the absence of WRC, while the half times of recovery for this component remains largely unchanged.

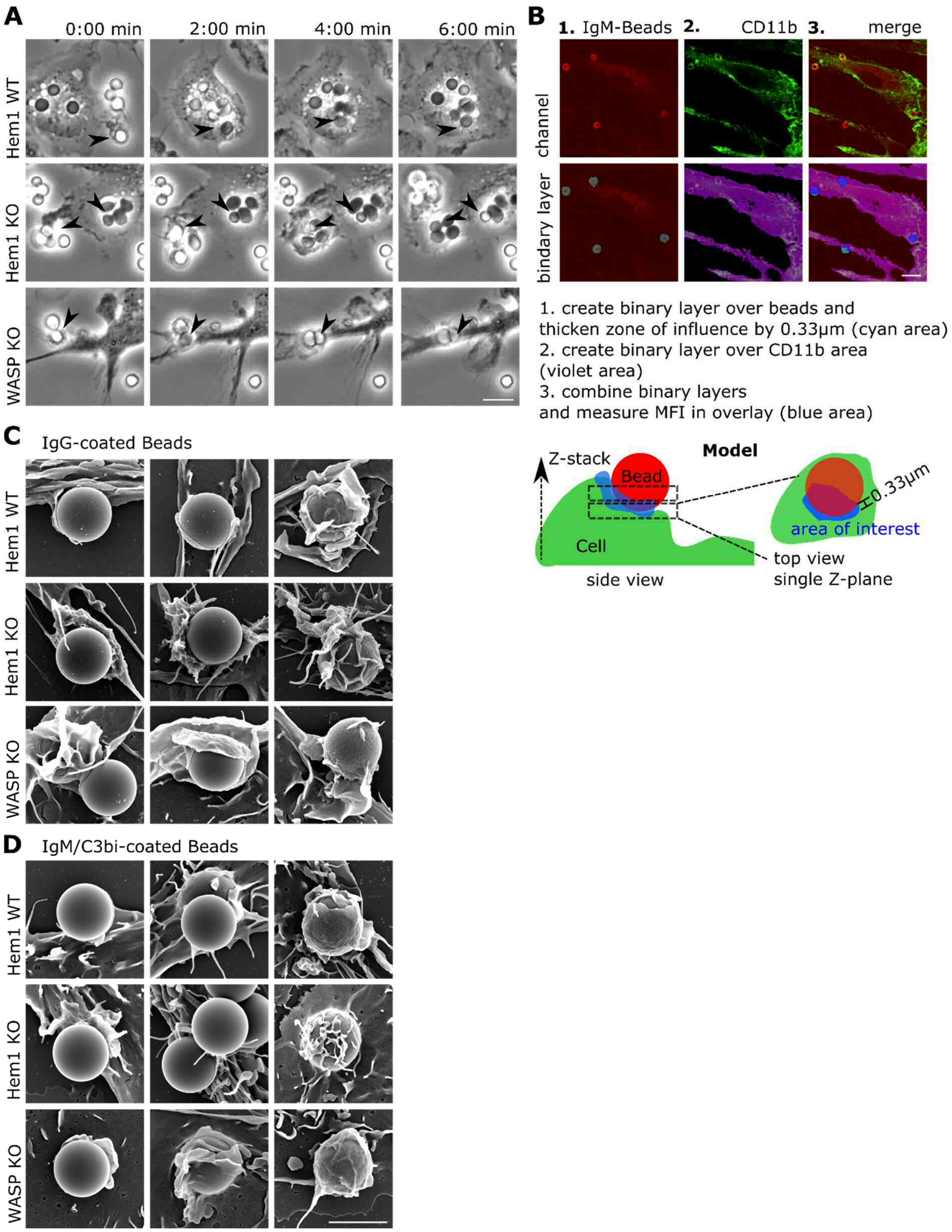


**Figure S4. FCγR- and CR3-dependent phagocytosis**

**(A)** Video microscopy images (phase contrast) of macrophages engaging IgG-opsonized SRBCs (arrowheads). SRBCs are first bright and then turn dark in phase contrast when internalized before being dis-integrated. Bar=5µm. **(B)** Illustration of CD11b quantification performed for Fig. 4F. Bar=10µm. **(C, D)** Additional examples of SEM images depicting different stages of internalization of macrophages phagocytosing either IgG- or IgM/C3bi-opsonized beads in subpanels C and D, respectively. Bead diameter: d=3µm, bar=3µm.

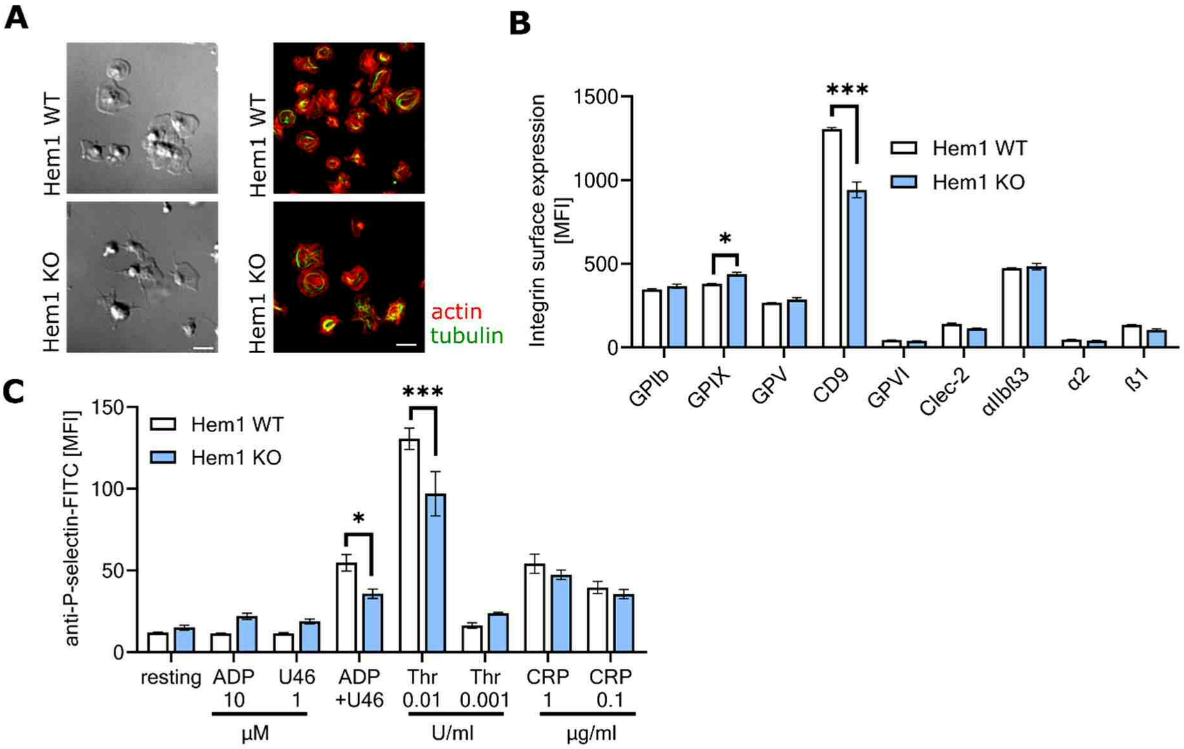


**Figure S5. Cytoskeleton staining and adhesion molecule surface patterns of Hem1 KO platelets**

**(A)** Left: additional DIC images of WT and Hem1 KO platelets. Right: Confocal images of the actin cytoskeleton (red) and of microtubules (green) from Hem1 KO and WT platelets allowed to spread for 30min, as in Fig. 5C. Note that lamellipodia-like structures can be detected in platelets from Hem1 KO mice, albeit at reduced frequency as compared to controls. Bar=5µm. **(B)** Mean fluorescence intensity of important platelet surface receptors, as determined by flow cytometry (n=5). Note that only CD9, the co-receptor of αIIβ3, is reduced in its surface prevalence. **(C)** Degranulation-dependent P-selectin exposure in response to indicated agonists (n=4).

**Supplemental movie legends**

**Movie S1A. Random Migration**

Phase contrast time-lapse microscopy illustrating the distinct morphologies of Hem1 WT, Hem1 KO and WASP KO macrophages during random migration. Note the spiky appearance and aberrant migration of Hem1 KO macrophages. Time is given in hh:mm.

**Movie S1B. Cremaster**

Movie displaying a 3D reconstruction of confocal images from cremaster tissue of male Hem1 WT, Hem1 KO and WASP KO mice. Myeloid cells are stained with CD11b (green) and endothelial cells are stained with CD31 (red). Note the strong accumulation of CD11b cells in and around blood vessels in Hem1 KO mice.

**Movie S2. Detachment of macrophages**

Macrophages of genotype as indicated were first stained with the viability marker Ca-AM (green) and then treated with 1mM EDTA in PBS to induce detachment. Note the progressive rounding of Hem1 WT and WASP KO macrophages, which is delayed in Hem1 KO macrophages. Time is hh:mm.

**Movie S3A. Detachment of B16 cells**

B16 cells were first stained with the viability marker Ca-AM (green) and then treated with 1mM EDTA in PBS to induce detachment. Note the progressive rounding of B16-F1 cells, which is delayed in Hem1/Nap1 and Sra-1/PIR121 double knockout clones (left panel). The right panel shows the analysis mask (violet border) used to measure the eccentricity of cells. Time is hh:mm.

**Movie S3B. FRAP of EGFP-tagged paxillin**

Movies of B16-F1 (left) and Sra-1/PIR121 KO#3 (right) cells expressing EGFP-tagged paxillin. Cells were imaged every 5 seconds by epifluorescence microscopy before and after local bleaching of individual focal adhesions (arrowheads) with a 405nm UV-laser. Note that movies were transiently frozen immediately before bleaching, to best illustrate recovery of fluorescence following individual bleaching events. Time is mm:ss, bar=5 µm.

**Movie S3C. FRAP of EGFP-tagged VASP**

Movies of cells expressing EGFP-tagged VASP and illustrating fluorescence recovery after photobleaching of individual focal adhesions marked by arrowheads, essentially performed as described for paxillin in Movie S3B. Time is mm:ss, bar=5 µm

**Movie S4A. Phagocytosis of phrodo™ Zymosan particles**

Overlay of phase contrast and epifluorescence time-lapse images of Hem1 WT, Hem1 KO and WASP KO macrophages phagocytosing phrodo™ Zymosan particles (green). Note the increase of green fluorescence over time in Hem1 WT, and the delayed response observed in Hem1 KO and WASP KO macrophages. Time is hh:mm, bar=5µm

**Movie S4B. Phagocytosis of IgG-SRBCs**

Phase contrast time-lapse microscopy illustrating phagocytosis of IgG-opsonized SRBCs by Hem1 WT, Hem1 KO and WASP KO macrophages. Accomplished phagocytosis of SRBCs darkens their appearance in phase contrast imaging from white to dark grey/black. Note that SRBCs are frequently not completely phagocytosed by Hem1 KO macrophages (red arrow). Time is hh:mm:ss.
